# Environmental legacy effects impact maize growth and microbiome assembly under drought stress

**DOI:** 10.1101/2023.04.11.536405

**Authors:** Joel F. Swift, Matthew R. Kolp, Amanda Carmichael, Natalie E. Ford, Paige M. Hansen, Benjamin A. Sikes, Manuel Kleiner, Maggie R. Wagner

## Abstract

**Background and Aims:** As the climate changes, plants and their associated microbiomes face greater water limitation and increased frequency of drought. Historical environmental patterns can leave a legacy effect on soil and root-associated microbiomes, but the impact of this conditioning on future drought performance is poorly understood. Precipitation gradients provide a means to assess these legacy effects.

**Methods:** We collected soil microbiomes from four native prairies across a steep precipitation gradient in Kansas, USA. Seedlings of two *Zea mays* genotypes were inoculated with each soil microbiome in a factorial drought experiment. We investigated plant phenotypic and root microbiome responses to drought and modeled relationships between plant growth metrics and climatic conditions from the soil microbiome origin sites.

**Results:** Drought caused plants to accumulate shoot mass more slowly and achieve greater root/shoot mass ratios. Drought restructured the bacterial root-associated microbiome via depletion of Pseudomonadota and enrichment of Actinomycetota, whereas the fungal microbiome was largely unaffected. An environmental legacy effect on prairie soil microbiomes influenced plants’ drought responses: counterintuitively, prairie soil inocula from historically wetter locations increased shoot biomass under drought more than inocula from historically drier prairie soils.

**Conclusion:** We demonstrated links between soil microbiome legacy effects and plant performance under drought, suggesting that future drying climates may condition soils to negatively impact plant performance.

## Introduction

Increasing drought frequency due to global climate change will impact both soil and plant-associated microbiomes (Chiang et al. 2021; Balting et al. 2021). Drought directly limits microbial growth, mobility, and access to resources, but also alters plant host traits (e.g., root physiology, morphology, and exudate composition) that comprise the habitat available to root-associated microorganisms (Naylor and Coleman-Derr 2018). Historical environmental patterns and drought can leave lasting impacts, or legacy effects, on soil and root-associated microbiomes. Naturally-occurring precipitation gradients—which shape the biogeography of both plants (Field et al. 2009; Korell et al. 2021) and soil microorganisms (Bachar et al. 2010; Hawkes et al. 2011)—provide opportunities to study legacy effects of the environment on plant-microbe interactions. Historical patterns of water availability across gradients constrain the functional potential of soil microbiomes (Hawkes and Keitt 2015; Averill et al. 2016; Canarini et al. 2021); however, the implications for host plants, including crops, are not well understood.

Because plants recruit root microbes primarily from the soil (Zarraonaindia et al. 2015), we would expect drought-induced changes in the soil microbiome to be reflected in root-associated microbiomes. However, because many root-associated microbes are also sensitive to plant root traits, their responses to drought may also reflect drought-induced changes in the host phenotype. Both soil- and root-associated microbiomes under drought exhibit consistent enrichment of Gram-positive bacteria and depletion of Gram-negative bacteria (Acosta-Martínez et al. 2014; Fuchslueger et al. 2016; Naylor et al. 2017). Within those categories, however, some bacterial phyla respond to drought differently in soil versus roots (Naylor and Coleman-Derr 2018). For example, enrichment of Bacillota (formerly Firmicutes) and Actinomycetota (formerly Actinobacteria) during drought are less pronounced in soil than in root-associated microbiomes, suggesting that these phyla respond to drought-induced changes in root physiology (Naylor et al. 2017; Fitzpatrick et al. 2018). Furthermore, the presence of a host plant can influence the drought responses of the taxa that comprise each phylum (Santos-Medellín et al. 2017). Overall, drought impacts both soil- and root-associated microbiomes, but in different ways, owing in part to the influence of the plant phenotype.

Changes in soil- and root-associated microbiomes caused by historical precipitation may affect plant fitness under drought. Drought-induced restructuring of root-associated microbiomes is hypothesized to be a conserved mechanism that supports survival of both the plant host and its symbionts (Naylor et al. 2017; Xu et al. 2018; Xu and Coleman-Derr 2019). For example, studies have shown that plant fitness is increased when a soil microbiome’s historic moisture conditions match contemporary experimental conditions (Lau and Lennon 2012; Munoz-Ucros et al. 2022), with this effect observed more commonly with co-adapted plants and soils that historically share a habitat (Lau et al. 2017; Remke et al. 2021; de Vries et al. 2023). Additional research suggests that exposure to short-term drought creates a microbial legacy effect that moderates drought stress in plants (Kaisermann et al. 2017; De Long et al. 2019) and strengthens positive plant-soil feedbacks, although many such patterns were species-specific (Buchenau et al. 2022). Thus, how historical precipitation patterns condition soil- and root-associated microbiomes and their relevance to plant fitness under drought stress is an open question.

To address this open question about legacy effects on microbiome function, we designed an experiment to assess plant fitness under a contemporary drought when inoculated with soil microbiomes from different historic precipitation regimes. Specifically, we collected soil microbiomes from native prairie remnants spanning an east-west precipitation gradient in Kansas (Figure 1). This precipitation gradient is coupled with multiple climatic metrics, with the environment becoming increasingly arid from east to west resulting in lower soil moisture and increased reference evapotranspiration (Figure 1C-E). We inoculated *Zea mays* L. (maize) seedlings with these soil microbiomes and measured plant growth during our experimental drought. Our goals were to: **1**) characterize plant phenotype and microbiome responses to drought, **2**) test how soil microbiome histories affected maize growth, and **3**) identify microbial taxa that were associated with plant growth. Our results suggest that both historical and experimental drought shape soil microbiomes’ interactions with maize.

**Figure 1.**
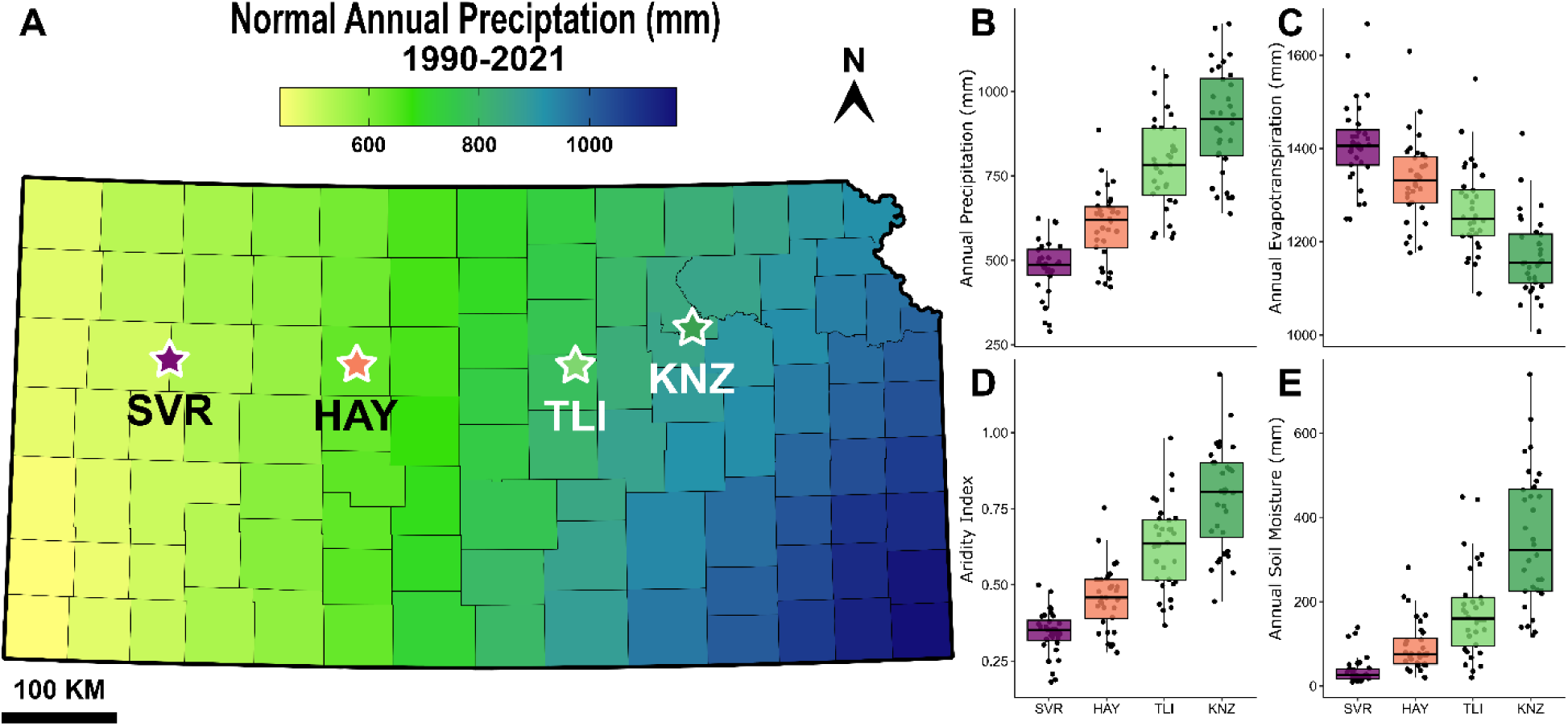
Kansas precipitation gradient. **A)** Soil samples were collected from four prairies in Kansas along an increasing precipitation gradient. Locations from east to west, Smoky Valley Ranch prairie (SVR; latitude 38.8665, longitude −100.9951), Hayes prairie (HAY; 38.8355, - 99.3033), The Land Institute prairie (TLI; 38.9698, −97.4690), and Konza native prairie (KNZ; 39.1056, −96.6099). Boxplots to the right of the map depict annual climatic metrics extracted from the TerraClimate database for each collection site from 1990-2021: **B)** precipitation, **C)** reference evapotranspiration, **D)** aridity index, and **E)** soil moisture. Boxplot hinges represent the 1^st^ and 3^rd^ quartiles; whiskers represent 1.5 times the interquartile range.

## Material and Methods

### Soil inoculum sources and preparation

Soil samples were collected in May 2019 from four prairie sites across a steep precipitation gradient in Kansas, USA (Figure 1). Prairies were selected at the same latitude and relative elevation such that they have very similar temperature profiles throughout the year. Western prairie sites (SVR and HAY) are predominantly short- to mixed-grass prairies (e.g., *Bouteloua* spp.), while eastern sites (TLI and KNZ) are dominated by tallgrass prairie species (e.g., *Andropogon gerardi*, *Schizachyrium scoparium*, *Panicum virgatum*). All collection sites were high-quality remnant prairies that have never experienced tillage or irrigation, which are strong perturbations that may obscure environmental legacy effects. Soil was collected from five evenly-spaced sub-samples per site, sampling to an approximate depth of 60 cm. Soil sub-samples were homogenized and sieved to remove rocks and plant material and then stored in plastic bags at 4°C until use, approximately four months after collection.

Inocula were prepared by mixing 20 g of each soil into 100 mL 1x phosphate-buffered saline (PBS) and 0.0001% Triton X-100. After settling, the soil slurries were filtered through Miracloth (22-25 µm pore size; Calbiochem, San Diego, CA, USA) and then centrifuged for 30 minutes (3000 x g). After discarding the supernatant, we resuspended the microbial pellets in 20 mL 1x PBS and kept them shaking for aeration until use, later that day. Each final inoculum consisted of 10 mL of the resuspended pellet in 1 L 0.5X Murashige and Skoog basal salt mixture (MS; Sigma-Aldrich, Darmstadt, Germany). A control treatment was made by adding 10 mL sterile 1x PBS into 1 L 0.5X MS.

### Planting, inoculation, and greenhouse setup

To increase the generalizability of our results, two genotypes of *Zea mays* L. (maize) were used: B73 and Mo17. These genotypes have been shown to vary in their growth response to beneficial endophytes (Schultz et al. 2022) and so may differ in response based on microbiome origin. In a laminar flow hood, seeds were surface sterilized by submerging in 70% ethanol, then 5% NaClO, followed by three rinses with sterile water. To avoid confounding effects of soil physicochemical properties, autoclaved calcined clay (“Pro’s Choice Rapid Dry”; Oil-Dri Corporation, Chicago, IL) was used as the growth medium. Seeds were planted one inch deep in 107mL “cone-tainers” (Stuewe & Sons Inc., Tangent, OR, USA) with a sterilized cotton ball pushed to the bottom to prevent clay from passing through drainage holes. Approximately 100mL of sterilized calcined clay was used per cone-tainer. The day after planting, cone-tainers were assigned randomly to treatments, inoculated with 25 mL of one of the four soil inocula (or the sterile control), and placed two inches apart in randomized complete blocks. Overall, each combination of treatments (drought treatment × soil inocula × genotype) was replicated 29 to 31 times across 27 randomized blocks (Figure S1A). Plants were grown in a temperature-controlled greenhouse (23/20 °C day/night, respectively) at the University of Kansas (Lawrence, KS, USA) arranged under growth lights (250 W Sun Systems Compact Cool fluorescent lamps) to supplement natural light for 13 hours/day during the experiment (January-March 2020).

### Monitoring and harvesting plants

All plants received 30 mL of reverse osmosis water twice weekly until emergence. After emergence, "well-watered” plants continued twice-weekly watering while “droughted” plants were watered every 8-12 days. Plant height was measured weekly to the nearest 0.25 cm on the tallest fully expanded leaf held parallel to the stem. Plants were sequentially harvested to measure biomass at one of three time points during the experiment: 68 plants on day 24 and day 39, and the remainder (n=224) on day 50 (Figure S1B). Calcined clay was removed from roots by hand. Shoot and root fractions were separated and oven-dried at 65 °C to a constant weight. Before drying, one nodal root (from the uppermost soil-borne root whorl) per plant was collected and stored at −80 °C until DNA extraction.

### Phenotypic analysis

All statistical analysis for phenotypic and microbiome data (see below) was conducted in R v3.6 (R Core Team 2021). Differences among group means were assessed using analysis of variance (ANOVA) with Type III sums of squares in the car v3.0.13 (Fox and Weisberg 2019) or lmertest v3.1.3 packages (Kuznetsova et al. 2017) with post-hoc comparisons made using the emmeans package v1.7.4.1 (Lenth et al. 2020). Assumptions of all models were checked by inspecting model residuals (Harrison et al. 2018). Dry shoot and root mass were converted to accumulation rates (g/day) to account for the variable number of growing days plants experienced post-emergence. The lme4 package v1.1.29 (Bates et al. 2015) was used to fit a generalized linear mixed model for emergence (binary) and linear mixed models for other growth responses (continuous). Models used the following formula (*growth rate ∼ Inoculum* × *Treatment* × *Genotype*) with all possible interactions included. Experimental block was treated as a random effect with time point included as an additional random effect in the root/shoot ratio model. Root/shoot ratio was natural log-transformed, and both shoot and root mass accumulation rates were square-root-transformed to meet the assumptions of ANOVA.

Models were also fit using climate metrics to test for linear relationships between plant growth and climatic conditions from the soil microbiome origin sites. Monthly precipitation, reference evapotranspiration (grass reference crop), Palmer’s Drought Severity Index (PDSI), and soil moisture values were extracted from TerraClimate database (Abatzoglou et al. 2018), selecting the closest 4×4 km grid cell to each site. Annual normals (30-year averages; Figures 1B-D and S2) were calculated for each metric, PDSI was calculated as a monthly average to reduce autocorrelation, and an aridity index was calculated by dividing normal annual precipitation by evapotranspiration.

### DNA extraction and library preparation

Nodal roots were freeze-dried for 48 hours (FreeZone lyophilizer; Labconco, Kansas City, MO, USA) and ground to powder with a HT Lysing Homogenizer (OHAUS, Parsippany, NJ, USA). Each ground root sample was mixed into 800 µL lysis buffer (1M Tris pH=8.0; 0.5 mM NaCl; 10 mM EDTA) and transferred to 96-well plates containing 500 µL of 2 mm garnet rocks per well (Biospec Products, Bartlesville, OK, USA), after which DNA purification proceeded via bead-beating and chemical lysis as previously described (Wagner et al. 2020).

Amplicon sequencing libraries were prepared using established protocols and PCR conditions (Wagner et al. 2020). Briefly, we targeted the V4 region of the 16S SSU rRNA gene for bacteria/archaea using 515F-806R primers (Apprill et al. 2015; Parada et al. 2016) and the ITS1 region for fungi using ITS1f and ITS2 primers (White et al. 1990; Gardes and Bruns 1993). Both amplicons were amplified separately using an initial PCR and then barcoded via a second index PCR. Amplicons were sequenced (2×250 bp) on a NovaSeq 6000 SP flow cell (Illumina, San Diego, CA, USA). Each “batch” of 96 samples (n=6) remained together throughout DNA extraction, amplification, and sequencing; individual samples were randomized across batches.

To confirm that we inoculated the experimental plants with realistic microbial communities, we also sequenced the original soils and freshly derived inocula. From homogenized soil collected from each site, we created eight technical replicates which we used to test whether our protocol for creating microbial slurries preserved the original community composition. DNA from each soil replicate and from freshly generated inocula was extracted in December 2022 using the same protocol that we used for roots, except omitting the lyophilization step and placing 250 mg soil or 800 µL inoculum directly into the garnet plate. From there, library preparation and sequencing proceeded as described above for roots.

### Bioinformatic processing

Sequence processing and quality filtering were conducted using an established workflow (Wagner et al. 2020). Briefly, Cutadapt v2.3 (Martin 2011) was used to remove primers and demultiplex reads into 16S and ITS sequence files. DADA2 v1.14.1 (Callahan et al. 2016) was used to denoise reads, merge paired-end sequences, dereplicate, and remove chimeric sequences. The forward and reverse 16S reads were truncated to 220 and 170bp, respectively and filtered based on the maximum number of expected errors of eight and nine bp per read, respectively. ITS reads were processed using DADA2 in an identical manner, except that reads were not truncated to preserve biologically relevant length variation. Forward and reverse 16S and ITS reads were then used separately to estimate sequencing error rates for denoising prior to dereplication of reads into Amplicon Sequence Variants (ASVs) and merging of pair-end reads. Taxonomy was assigned to the resulting ASVs by a naïve Bayesian classifier (Wang et al. 2007) trained on the 11.5 release of the Ribosomal Database Project (Callahan 2017) for bacteria and the 01.12.2017 release of the UNITE database (UNITE Community 2017) for fungi. Phyloseq v1.40.0 (McMurdie and Holmes 2013) was used to remove host sequences (chloroplast and mitochondria; 0.35% of 16S reads) and ASVs unassigned to a kingdom. Chao1 richness and diversity indices (Shannon and Inverse Simpson) were calculated before the removal of ASVs not observed ≥25 times in ≥5 samples and samples with <400 reads (200 bacterial and 133 fungal samples); 98.0% and 94.5% of reads remained after host filtering and thresholding (bacterial and fungal reads, respectively). Finally, ASV counts were normalized via a centered-log-ratio (CLR) transformation using the ALDEx2 package v1.28.1 (Fernandes et al. 2014; Gloor et al. 2017). Due to low replication and low sequencing depth at the earlier timepoints, we analyzed only microbiome samples from the last time point. This retained 131 bacterial (mean read depth 123,554 ± SE 18,276) and 92 fungal samples (41,081 ± 10,129).

### Microbiome Analysis

To test the similarity of inocula to their soils of origin, a linear model was fit for each alpha diversity metric with collection site, substrate (soil *vs.* inoculum), and sequencing depth (Z-transformed) as main effects. Community composition was assessed by permutational multivariate analysis of variance (PERMANOVA) and constrained partial redundancy analysis (RDA). For PERMANOVA, the *adonis2()* function from the vegan package v2.6.2 (Oksanen et al. 2019) was used to model the matrix of between-sample Aitchison distances (Gloor et al. 2017) with collection site and substrate as main effects. The RDA was fit with the same model, removing variance attributed to ln(Sequencing Depth), and visualized using an ordination plot.

To investigate root microbiome responses, a model was fit for each alpha diversity metric (*Metric ∼ Soil Inoculum* × *Drought Treatment* × *Genotype*), with *Block* and *Batch* as random effects. The CLR-transformed abundance of each family was modeled to test responses to experimental factors. Across all models and where appropriate, p-values were adjusted to correct for multiple comparisons using a Benjamini-Hochberg correction (Benjamini and Hochberg 1995). To explore changes in community composition, a PERMANOVA model was fit (*Aitchison distance matrix ∼ Soil Inoculum* × *Drought Treatment* × *Genotype*) with permutations constrained by *Batch.* Main effects were tested for homogeneity of dispersions using the vegan package. Pairwise dispersions among factor levels were assessed using a permutation test with 999 permutations constrained by *Batch.* Constrained partial redundancy analysis of the Aitchison distance matrix was conducted to remove variance attributed to *Batch* and *ln(Sequencing Depth)* and to visualize the variance in microbial composition explained by experimental factors.

The machine-learning package caret v6.0.92 (Kuhn 2008) with ranger’s v0.14.1 (Wright and Ziegler 2017) implementation of the random forest algorithm was used to assess how accurately roots could be classified as droughted or well-watered based on microbiome community data (bacterial and fungal, separately). Samples were randomly split into a training set (80%) and a test set (20%), with the test set withheld from the model, and used to determine the final model’s performance. Hyperparameters were optimized using a grid search over minimum node size (1, 5, 10) and the number of features available at each node (10-100% of the ASVs). For each combination in the grid, classifier accuracy was used to assess performance on out-of-bag samples with 10-fold cross validation. Final predictions were made with the trained model on the withheld test set and visualized using confusion matrices. The contribution of each ASV to classification accuracy was assessed using permuted importance: mean decrease in accuracy (MDA) values were calculated for each ASV across the 10-fold cross validation. The benefits of this strategy are twofold, in that we can assess whether drought induced a predictable shift in the microbiome as a whole and identify individual ASVs with the greatest differential abundance as a result of drought.

### Connecting plant growth to microbial abundances

Linear models were used to test associations between taxon abundances and plant growth. To focus on the phenotypic variation that could not be explained by Genotype, Soil Inoculum, Block and Batch effects, we extracted residuals from a model that contained these terms and used them as the response variable for a second model (*Growth Response_residual_ ∼ Taxon abundance × Drought Treatment*). The *Taxon abundance* coefficient described the association of the taxon (CLR-transformed counts of bacterial and fungal families) with plant growth rate; the interaction term described whether the taxon’s effect on plant growth differed under droughted vs. well-watered conditions. We chose family and order as the taxonomic levels for testing bacteria and fungi, respectively, as a conservative measure to limit the number of tests. P-values were adjusted by term to account for multiple testing.

## Results

### Soil and inocula community composition differs across collection sites

We assessed the microbial community composition of soils and their derived inocula from the four collection sites (n=8 samples per site). Bacterial, but not fungal, alpha diversity of soils differed across collection sites (Tables S1 and S2); bacterial diversity was highest in the westernmost prairie soil (Figure S3). Community composition of both bacterial and fungal microbiomes differed among sites (Table S3; R^2^=0.22-0.24; *p*-value<0.01). Relative abundances of bacterial phyla and fungal classes further showed site-specific patterns (Figures S5 and S6).

Relative to the original soils, we observed that inocula generally had lower richness and diversity, but this difference was non-significant for most metrics (Figure S4; Tables S1 and S2). The composition of both bacterial and fungal communities also shifted between soils and their derived inocula (Figure S5-6; Table S3); however, both soil and inocula samples clustered in ordination space by collection site (Figure 2). Thus, the derived inocula largely retained the differences among the original soil microbial communities.

**Figure 2.**
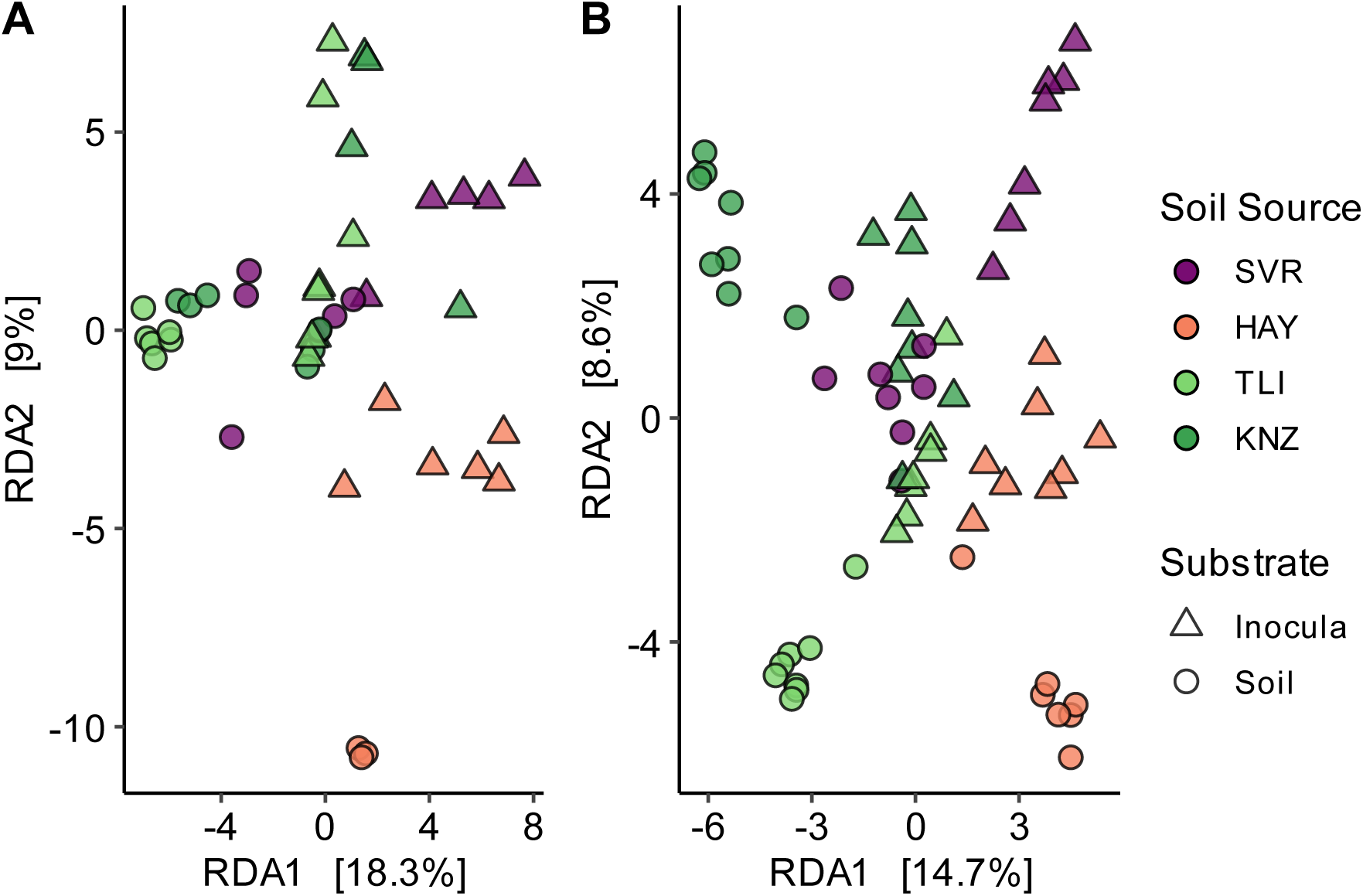
Soil community composition is strongly differentiated across the soil sources and substrates. Redundancy analysis for **A**) bacterial and **B**) fungal communities from soil sources used to derive soil inocula. Soil source and substrate are represented by differing colors or shapes, respectively.

### Plant growth rates vary across soil microbiomes, genotype, and drought treatment

We found no evidence that soil inocula impacted emergence rates (*p*-value = 0.19, Pearson’s χ^2^ test; Table S4), however, maize genotype B73 (81.3%) emerged more successfully than Mo17 (62.0%; *p*-value=0.05, Pearson’s χ^2^ test). Post-emergence, both drought treatment and maize genotype affected plant growth (Table 1). Drought decreased both shoot and root mass accumulation rates (Figure 3A-B; *p*-value=<0.01) and increased root/shoot ratio (Figure 3C; *p*-value<0.01). These traits also differed between genotypes, with Mo17 investing more into root growth than B73 (Figure S7; *p*-value=<0.01). Shoot mass accumulation rate and root/shoot ratio were both influenced by soil inoculum (Table 1; *p*-value=<0.01-0.02); this effect was driven by the ‘TLI’ inoculum which caused the slowest accumulation of shoot mass and the lowest root/shoot ratio. We observed no evidence of interactions between drought treatment, genotype, and soil inoculum on plant growth.

**Figure 3.**
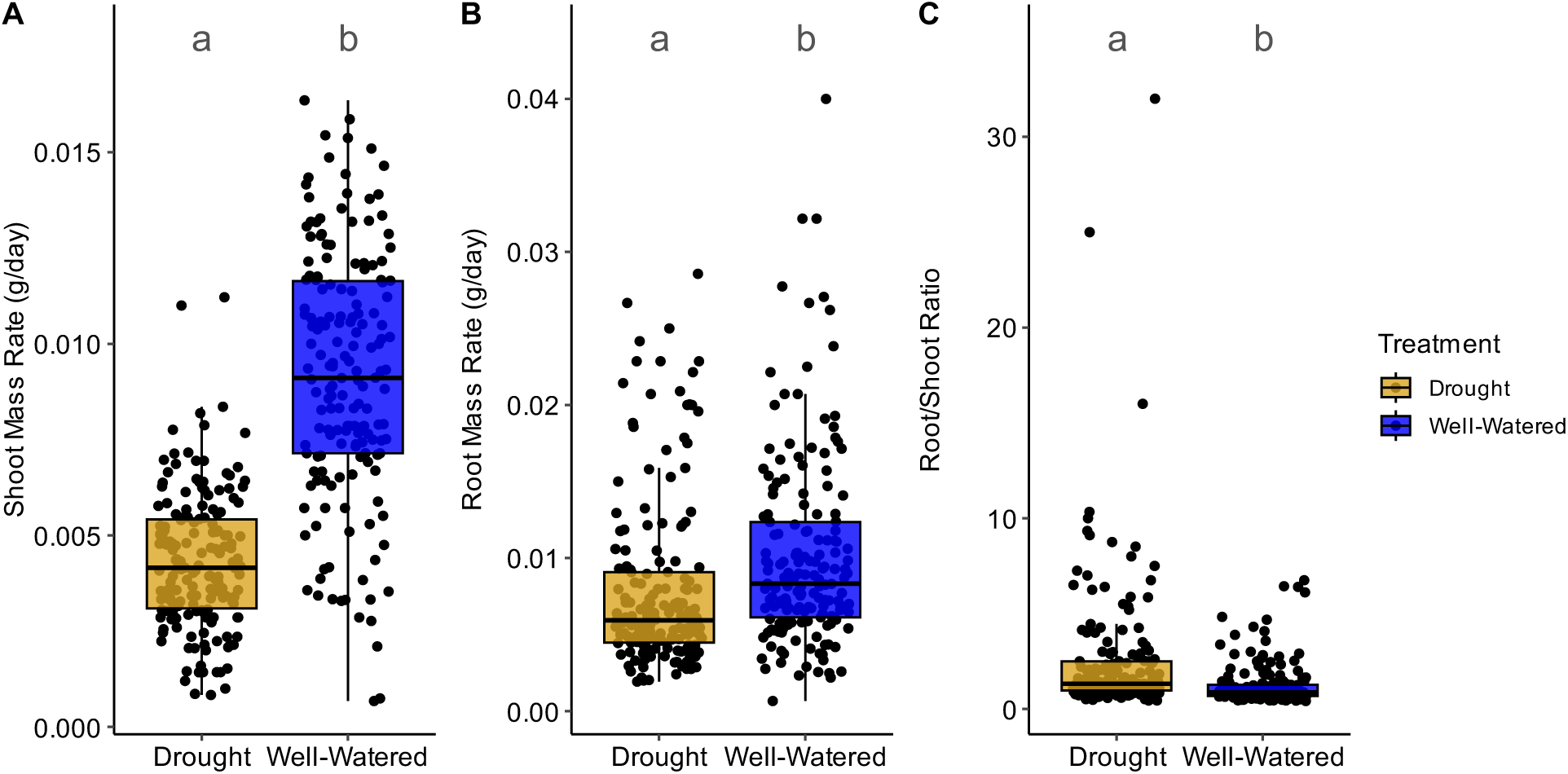
Shoot and root growth vary across drought treatment. The effect of drought treatment on **A)** shoot mass accumulation rate, **B)** root mass accumulation rate, and **C)** root/shoot ratio. Boxplot hinges represent the 1^st^ and 3^rd^ quartiles; whiskers represent 1.5 times the interquartile range; letters above box plots denote significant differences in estimated marginal means (*p*-value=<0.01; see table 1 for full ANOVA table).

**Table 1.** Type three analysis of variance (ANOVA) for plant growth.

| Factor | DF | sqrt(Shoot mass rate) |  | sqrt(Root Mass Rate) |  | log(Root/shoot Ratio) |  |
| --- | --- | --- | --- | --- | --- | --- | --- |
| | | $F$-value | $p$-value | $F$-value | $p$-value | $F$-value | $p$-value |
| **Drought Treatment (DT)** | 1 | 335.67 | **<0.01** | 16.96 | **<0.01** | 161.21 | **<0.01** |
| **Genotype (G)** | 1 | 33.56 | **<0.01** | 22.74 | **<0.01** | 110.11 | **<0.01** |
| **Soil Inoculum (SI)** | 3 | 5.08 | **<0.01** | 0.67 | 0.57 | 3.30 | **0.02** |
| **DT x G** | 1 | 0.08 | 0.77 | 0.93 | 0.34 | 0.12 | 0.73 |
| **DT x SI** | 3 | 2.25 | 0.08 | 1.73 | 0.16 | 0.63 | 0.59 |
| **G x SI** | 3 | 2.13 | 0.10 | 0.61 | 0.61 | 0.44 | 0.73 |
| **DT x G x SI** | 3 | 0.25 | 0.86 | 0.09 | 0.96 | 0.13 | 0.94 |

Bold values denote significant effects, $p$ -value $<0.05$

### Root microbiomes and community responses to drought differed between bacteria and fungi

We observed different patterns of richness and diversity for bacterial and fungal root microbiomes exposed to prairie inoculum and drought (Tables S5 and S6). Bacterial Chao1 richness varied across soil inocula (Table S5; *p*-value=0.03), with roots receiving the ‘KNZ’ inoculum exhibiting the highest richness. Drought increased Shannon and Inverse Simpson’s diversity of fungal communities (Table S6). We observed no difference in bacterial or fungal richness or diversity between the maize genotypes (Tables S5 and S6).

Root bacterial community composition was also structured by soil inoculum (Table 2; R^2^=0.05, *p*-value=<0.01, PERMANOVA) and drought treatment (Figure 4A; R^2^=0.01, *p*-value=0.01, PERMANOVA). We found no evidence that bacterial community dispersion differed among treatments, genotypes, or inocula (*p*-value>0.05, Multivariate Levene’s test). Redundancy analysis aligned with PERMANOVA results: soil inoculum and drought treatment explained the largest proportions of variance (6.09% and 1.57%; Figure S8 and 4B, respectively), while genotype explained <1% of variance (Figure S9). Among bacterial families, Sphingomonadaceae was responsive to drought treatment (*q*=7.1e^-4^), with this family decreasing from 1.11% relative abundance to 0.46% under drought conditions.

**Figure 4.**
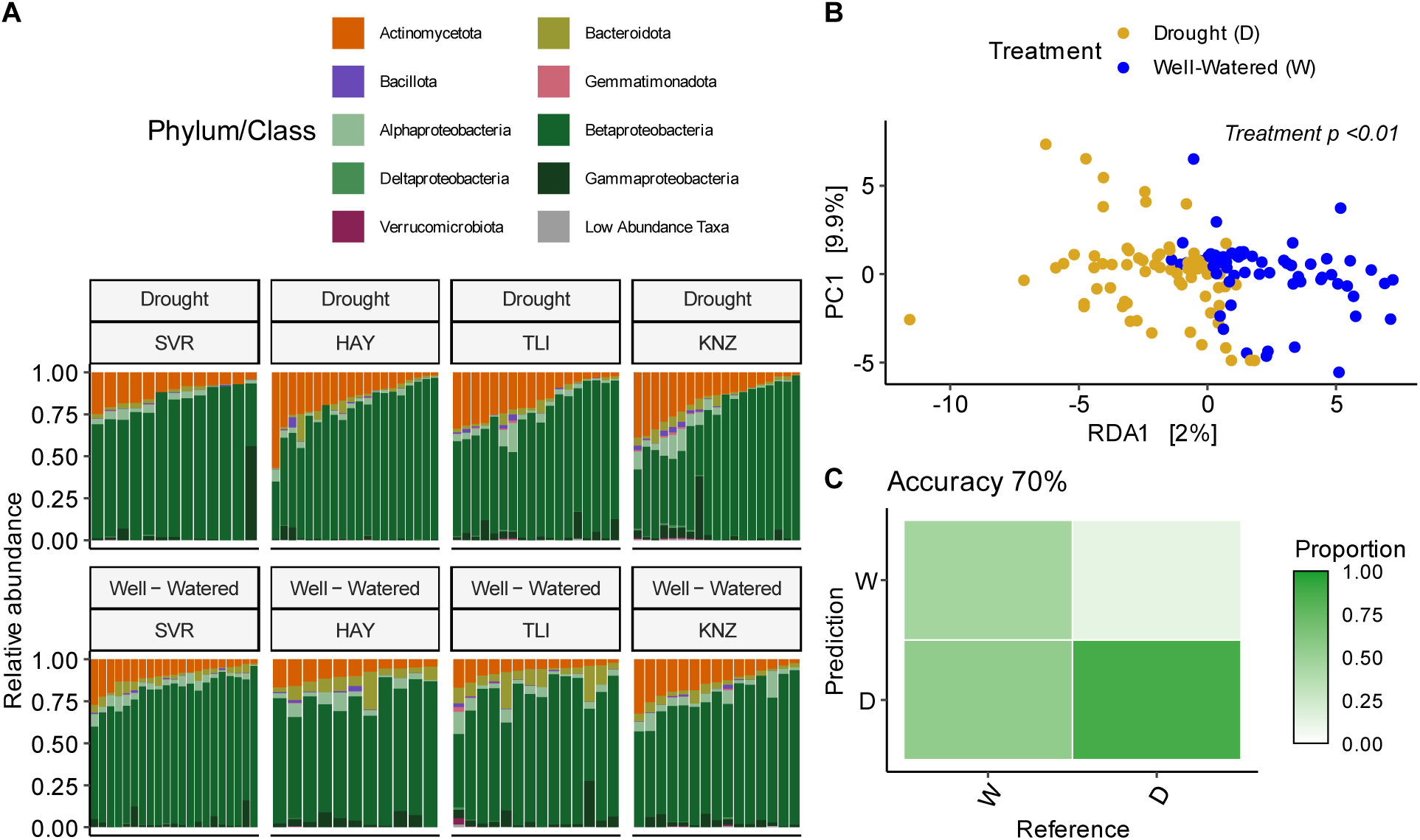
Bacterial community composition across drought treatments. **A)** Bacterial taxonomic barplots for drought-stressed and well-watered plants. Each site is represented by a facet with samples denoted by individual lines and the relative abundance (non CLR transformed data) of each phylum displayed by color. **B)** Redundancy analysis on bacterial communities constrained by drought treatment, conditioned on sequencing batch and ln-transformed usable reads per sample. **C)** Confusion matrix depicting the classification accuracy of a random forest classifier predicting drought treatment trained on bacterial communities. Above the confusion matrix balanced accuracy is shown; for additional classifier statistics, see Table S8.

**Table 2.** Permutational Multivariate ANOVA (PERMANOVA) table bacterial community composition of maize nodal roots.

| Factor | DF | SS | R<sup>2</sup> | F-value | p-value |
| --- | --- | --- | --- | --- | --- |
| Genotype (G) | 1 | 2687 | 0.01 | 0.88 | 0.47 |
| Soil Inoculum (SI) | 3 | 19091 | 0.05 | 2.10 | **<0.01** |
| Drought Treatment (DT) | 1 | 5484 | 0.01 | 1.81 | **0.01** |
| G x SI | 3 | 9849 | 0.02 | 1.08 | 0.47 |
| G x DT | 1 | 2289 | 0.01 | 0.75 | 0.72 |
| SI x DT | 3 | 10026 | 0.02 | 1.10 | 0.43 |
| G x SI x DT | 3 | 6352 | 0.02 | 0.70 | 0.95 |
| Residual | 115 | 349180 | 0.86 |  |  |

Bold values denote significant effects, $p$ -value<0.05

In contrast to bacteria, drought treatment did not alter relative abundances of any fungal families, nor overall root fungal community profiles (Figure 5A). Variance in fungal community composition was mostly due to initial soil inoculum source (Figure S10; Table 3; R^2^=0.09, *p*-value=<0.01). As observed for bacterial communities, neither drought treatment, maize genotype, nor soil inoculum source affected fungal community dispersion (*p*-value=>0.05, Multivariate Levene’s test). Drought and maize genotype each explained only a small portion of the variance in fungal community composition (Figure 5B and S11, respectively). Thus, while root-associated bacterial community composition responded to drought treatment, we observed no such trend for fungal community composition.

**Figure 5.**
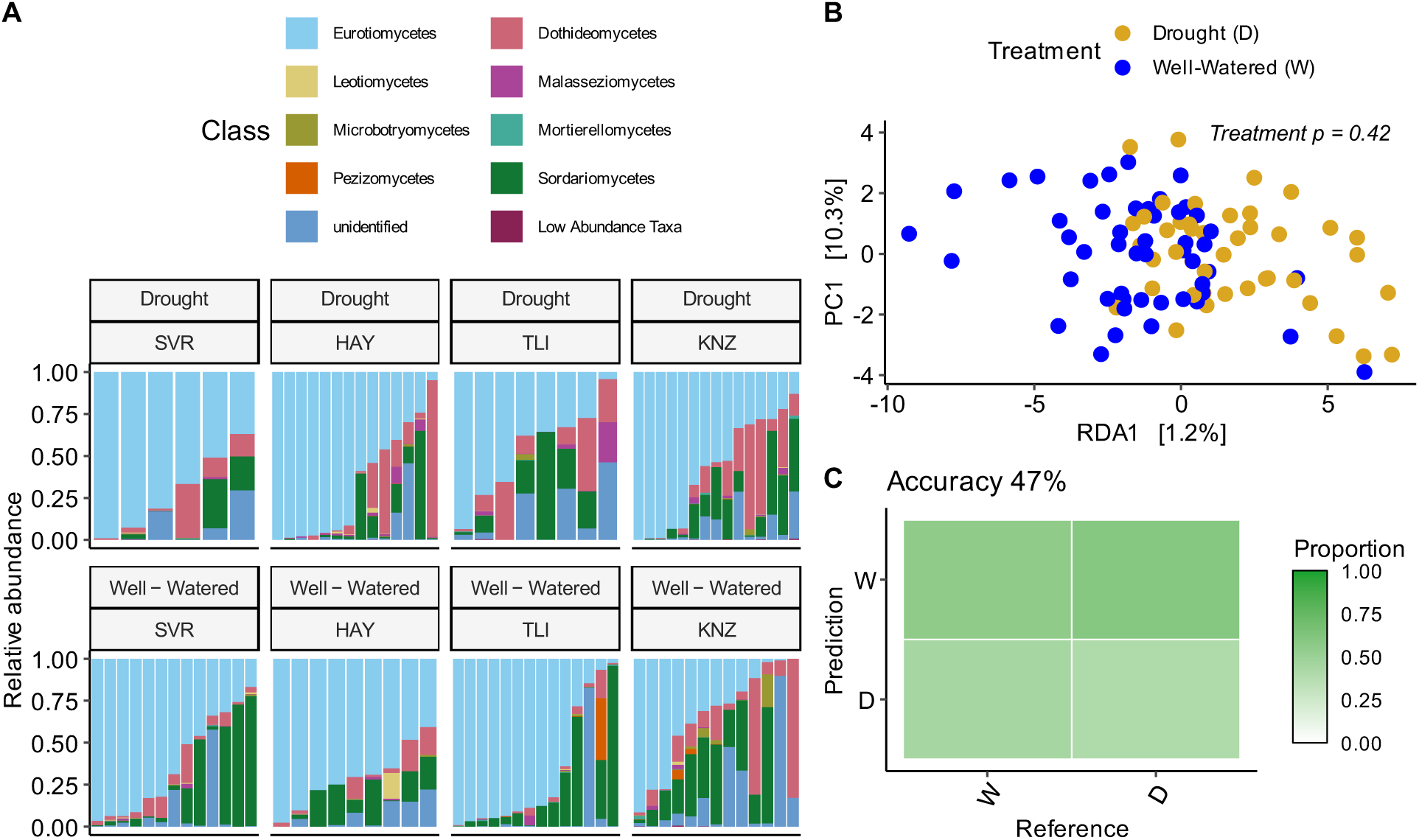
Fungal community composition across drought treatment. **A)** Fungal taxonomic barplots for drought-stressed and well-watered plants. Each site is represented by a facet with samples denoted by individual lines and the relative abundance (non CLR transformed data) of each phylum displayed by color. **B)** Redundancy analysis on fungal communities constrained by drought treatment, conditioned on sequencing batch and ln-transformed usable reads per sample. **C)** Confusion matrix depicting the classification accuracy of random forest a classifier predicting drought treatment trained on fungal communities. Above the confusion matrix balanced accuracy is shown; for additional classifier statistics, see Table S9.

**Table 3.** PERMANOVA table for fungal community composition of maize nodal roots.

| Factor | DF | SS | R<sup>2</sup> | F-value | p-value |
| --- | --- | --- | --- | --- | --- |
| Genotype (G) | 1 | 586 | 0.01 | 0.89 | 0.49 |
| Soil Inoculum (SI) | 3 | 5896 | 0.09 | 3.00 | **<0.01** |
| Drought Treatment (DT) | 1 | 541 | 0.01 | 0.83 | 0.63 |
| G x SI | 3 | 2181 | 0.03 | 1.11 | 0.60 |
| G x DT | 1 | 740 | 0.01 | 1.13 | 0.18 |
| SI x DT | 3 | 2335 | 0.04 | 1.19 | 0.17 |
| G x SI x DT | 3 | 1969 | 0.03 | 1.00 | 0.61 |
| Residual | 76 | 49822 | 0.78 |  |  |

Bold values denote significant effects, $p$ -value $<0.05$

### Identification of bacterial and fungal ASVs responding to drought

Our random forest models showed greater accuracy in predicting the drought treatment of a given sample when trained on bacterial communities than on fungal communities (Figures 4C and 5C). Three bacterial ASVs greatly aided the prediction of treatment (Figures 4C and S12-13): two from the family Sphingomonadaceae (Pseudomonadota) and one from the family Sphingobacteriaceae (Bacteroidota). These ASVs showed mean decrease in accuracy (MDA) values of 1.6%, 1.2%, and 1.2% when withheld from the model, respectively. In other words, these three bacterial ASVs contributed to ∼4% increase of classifier accuracy in predicting. All these bacterial ASVs were depleted in drought-stressed plants relative to well-watered plants: the Sphingomonadaceae ASVs decreased in relative abundance by an order of magnitude (0.37% and 0.31% to 0.03% and 0.04% relative abundance, respectively) and the Sphingobacteriaceae ASV responded similarly (0.9% to 0.1% relative abundance). We also detected two fungal ASVs that aided the prediction of treatment (Figure 5C and S14-15), from the families Aspergillaceae (1.1% MDA; Eurotiomycetes) and Sarocladiaceae (0.76% MDA; Sordariomycetes). These fungal ASVs showed different patterns in response to drought treatment: the Aspergillaceae ASV was enriched while the Sarocladiaceae ASV was depleted in drought-stressed plants.

### Connecting plant growth to bacterial and fungal communities

Soil inocula derived from collection sites across Kansas had only a small impact on plant growth (Table 1; Figure 6A and S15), with no single inoculum causing significantly improved growth over other inocula or the control. Examining the prairie sites spanning the east-west precipitation gradient, we detected an interaction between mean annual precipitation and drought treatment on shoot mass accumulation rate (Figure 6B; *p*-value=0.04). The shoot mass accumulation rate of well-watered plants was negatively associated with mean annual precipitation of the soil microbiome collection site (slope = −2.88e^-6^ g/day/mm precipitation) whereas droughted plants displayed a weak positive association (slope = 4.53e^-7^ g/day/mm precipitation). Put another way, counterintuitive to expectations, plants that experienced drought benefited from microbiomes from historically wetter soils, while the opposite was true for well-watered plants in the experiment (we discuss this striking pattern below). We observed that this pattern was consistent when substituting other climatic metrics (PDSI, aridity index, evapotranspiration, and soil moisture) for mean annual precipitation (Figure S17; *p*-values = 0.03, 0.06, 0.08, and 0.18, respectively).

**Figure 6.**
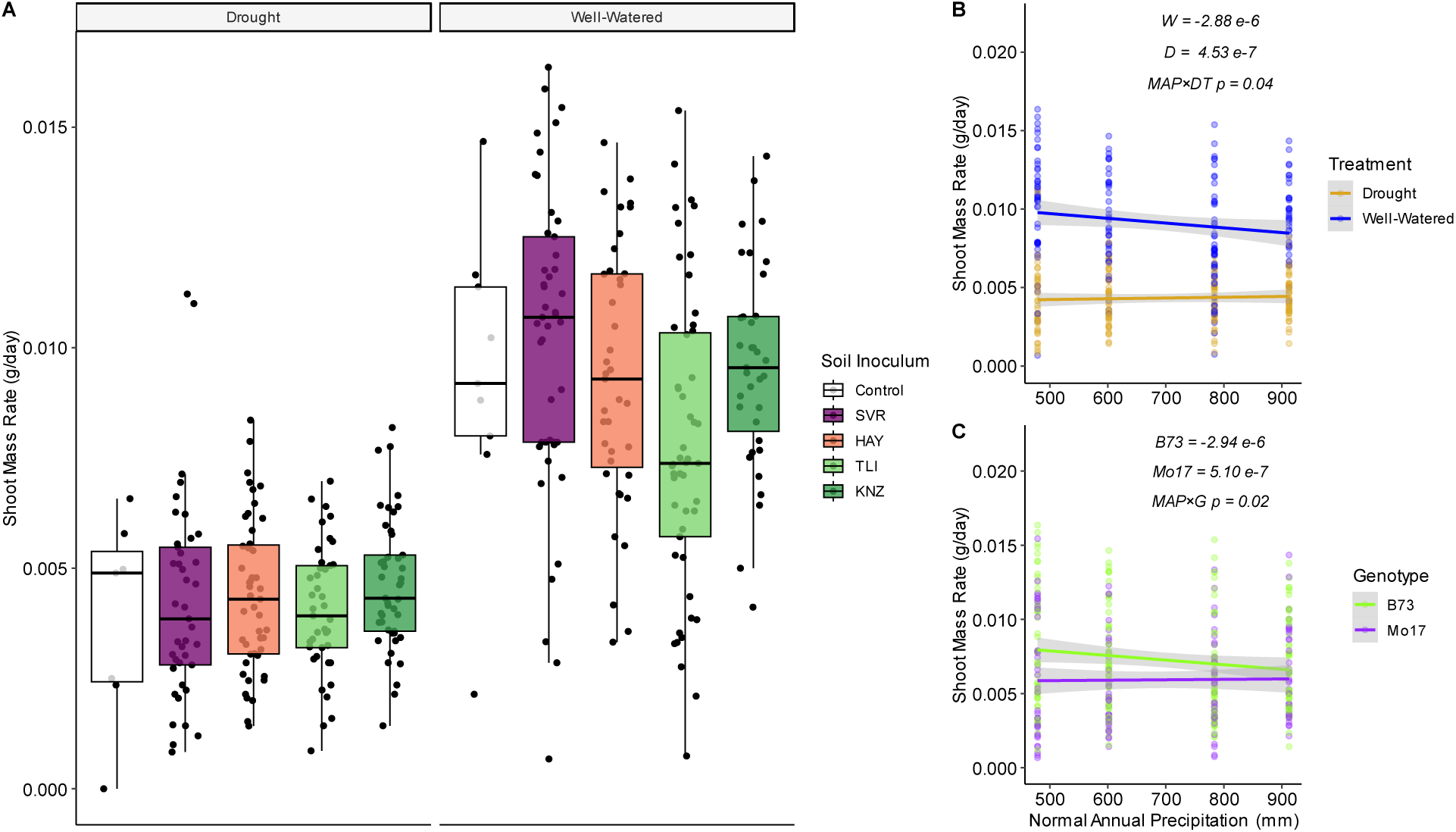
Shoot growth rate by soil inoculum. Boxplots are colored by soil inoculum for **A)** shoot mass accumulation rate with separate facets representing drought-stressed and well-watered plants. Boxplot hinges represent the 1^st^ and 3^rd^ quartiles; whiskers represent 1.5 times the interquartile range. **B-C)** shoot mass accumulation rate trend lines depicting significant interactions of mean annual precipitation (last climate normal 1991-2020) with drought treatment and genotype, respectively, for plants inoculated with prairie soils. For each regression line, grey shading represents the 95% confidence interval.

Notably, B73 and Mo17 exhibited contrasting associations with mean annual precipitation across the Kansas precipitation gradient (Figure 6C; *p*-value=0.02; Slope= −2.94e^-6^ and 5.10e^-7^, respectively); the same pattern was observed for other climatic metrics except for PDSI (Figure S17).

We identified bacterial and fungal taxa that were correlated with promotion or suppression of plant growth. Overall, 11 bacterial families showed a significant positive (3/11) or negative (8/11) association with shoot mass accumulation rate (Table 4). These represented only a few phyla (5 Actinomycetota, 4 Pseudomonadota, and 2 Bacillota) and were a mix of Gram-positive (7) and Gram-negative (5). One of the identified families, Streptococcaceae, interacted with drought treatment, indicating that its relationship with shoot mass accumulation was modulated by the amount of water the plant received. The abundance of Streptococcaceae was negatively correlated with shoot mass rate in well-watered plants but not in droughted plants (Figure S18). Relative to shoot mass accumulation rate, fewer bacterial families were associated with root mass accumulation rate (only Acetobacteraceae and Corynebacteriaceae, Table 4). This result may be due to variance in this trait decreasing over the course of the experiment as roots expanded to fill the entire container regardless of treatment or inoculum (Figure S19). Notably, Acetobacteraceae abundance was negatively associated with shoot mass accumulation rate but positively associated with root mass accumulation rate. We found that the families associated with root/shoot ratio, Sinobacteraceae and Sphingomonadaceae (Table 4), were a subset of families associated with shoot mass accumulation rate, suggesting that a change in shoot mass was responsible for this relationship. For fungi, only a single family, Sporidiobolaceae, was correlated with shoot mass accumulation rate (Table 4). Similar results were found at lower taxonomic levels for both bacteria and fungi (Table S10-11).

**Table 4.** Bacterial and fungal families associated with plant traits; square root transformed shoot mass accumulation rate (SMR), root mass accumulation rate (RMR), and root/shoot ratio (RSR).

| Taxon | Trait | Model Term | R | X | PVE<sub>Res</sub> | Rel. Abund. | q |
| --- | --- | --- | --- | --- | --- | --- | --- |
| Acetobacteraceae | SMR | Abundance | -1.25E-04 | -1.33E-02 | 6.67 | 0.02 | 2.76E-03 |
| Corynebacteriaceae | SMR | Abundance | -5.46E-04 | -1.29E-02 | 8.60 | 0.01 | 1.90E-04 |
| Dietziaceae | SMR | Abundance | -1.07E-03 | -1.31E-02 | 4.94 | 0.00 | 6.48E-03 |
| Lactobacillaceae | SMR | Abundance | 5.64E-04 | -1.46E-02 | 4.26 | 0.09 | 1.55E-02 |
| Microbacteriaceae | SMR | Abundance | -1.07E-05 | -1.32E-02 | 5.27 | 1.23 | 6.48E-03 |
| Micrococcaceae | SMR | Abundance | -8.43E-06 | -1.31E-02 | 3.84 | 8.47 | 2.46E-02 |
| Nocardiaceae | SMR | Abundance | -3.71E-04 | -1.33E-02 | 3.58 | 0.02 | 2.75E-02 |
| Rhodocyclaceae | SMR | Abundance | -4.92E-04 | -1.14E-02 | 3.51 | 0.06 | 2.75E-02 |
| Sinobacteraceae | SMR | Abundance | 2.12E-04 | -1.38E-02 | 3.70 | 0.05 | 2.07E-02 |
| Sphingomonadaceae | SMR | Abundance | -1.03E-04 | -1.24E-02 | 5.31 | 1.13 | 6.48E-03 |
| Streptococcaceae* | SMR | Abundance | 1.38E-04 | -1.34E-02 | 5.50 | 0.01 | 4.10E-03 |
| Acetobacteraceae | RMR | Abundance | 2.32E-05 | -5.94E-03 | 8.15 | 0.02 | 1.41E-02 |
| Corynebacteriaceae | RMR | Abundance | -2.65E-04 | -5.74E-03 | 6.29 | 0.01 | 3.39E-02 |
| Polyangiaceae | RSR | Abundance | 2.19E-02 | 1.37E-01 | 8.01 | 0.08 | 1.81E-02 |
| Sinobacteraceae | RSR | Abundance | -1.48E-03 | 1.67E-01 | 5.8 | 0.05 | 2.96E-02 |
| Sphingomonadaceae | RSR | Abundance | 6.96E-04 | 1.58E-01 | 7.44 | 1.13 | 1.81E-02 |
| Sporidiobolaceae | SMR | Abundance | 3.33E-04 | -1.24E-02 | 6.17 | 0.48 | 2.95E-02 |

\*Streptococcaceae shows a significant interaction between the family abundance and drought treatment, presented in Figure S11.

## Discussion

### Plant and microbiome responses to drought

Understanding plant-microbiome interactions under drought stress is critical to safeguard agricultural systems in the face of global climate change. In our experiment, maize displayed phenotypic tradeoffs in response to drought. Drought-treated plants increased root growth relative to shoot growth, resulting in shorter plants over the course of the experiment. This supports our understanding of plant responses to drought, as plants prioritize root over shoot growth (Sharp and Davies 1979; Lynch et al. 2014), favoring deeper and often denser root systems in order to obtain sufficient water (Zhan et al. 2015; Gao and Lynch 2016; Sebastian et al. 2016). The maize genotypes, B73 and Mo17, showed significant differences between root/shoot allocation patterns, but in general both genotypes responded to drought in a similar manner. These phenotypic changes in root growth, along with direct effects of water limitation in the soil, contributed to the responses of the root microbiomes in our study.

Drought altered the bacterial communities of maize roots, although the magnitude of compositional change within our study was small (R^2^=0.01) relative to other studies (R^2^=0.04-0.10) (Santos-Medellín et al. 2017; Naylor et al. 2017; Fitzpatrick et al. 2018; Xu et al. 2018). Similar to previous studies (Xu and Coleman-Derr 2019), we observed a depletion of Gram-negative and an enrichment of Gram-positive bacteria in response to drought, likely relating to the desiccation resistance provided by the thick peptidoglycan cell wall of Gram-positive bacteria (Manzoni et al. 2012). Enrichment of Actinomycetota during drought tends to be more pronounced in root-associated microbiomes than in soil, suggesting that shifts in plant root physiology also play a role (Naylor and Coleman-Derr 2018; Xu and Coleman-Derr 2019); however, the specific root traits driving this effect are unknown. During drought, root exudate production is generally maintained or increased (Henry et al. 2007; Preece and Peñuelas 2016; Karlowsky et al. 2018), while the exudate composition is typically altered (Svenningsson et al. 1990; Canarini et al. 2016; Calvo et al. 2017) which can contribute to the root microbiome drought response (Zhalnina et al. 2018). In *Sorghum*, for instance, drought-induced changes in root exudate composition and iron homeostasis contribute to Actinomycetota enrichment in the rhizosphere (Xu et al. 2018, 2021).

Fungal communities of maize roots were less sensitive to drought relative to their bacterial neighbors. While drought increased root fungal diversity in the plants inoculated with the TLI prairie soil inoculum, it did not significantly affect overall fungal community composition. Machine learning classification by drought treatment further illustrated this difference: when trained on bacteria, we achieved 70% balanced accuracy, while for fungi, we achieved only 47% accuracy. This discrepancy indicates that bacterial communities responded in a predictable manner to drought, whereas fungal communities were less predictable between well-watered and drought plants. This cross-kingdom difference agrees with previous reports, as both free living and root-associated fungal communities tend to tolerate desiccation (de Vries et al. 2012; Furze et al. 2017). The observed difference in survival between bacteria and fungi under drought may relate to both the vegetative and dormant growth forms of fungi: compared to bacterial cells, fungal hyphae comprise most vegetative growth and can cross air-gaps within dry soil in search of resources (Griffin 1985; Manzoni et al. 2012), and dormant spores are generally desiccation-resistant (Wyatt et al. 2013).

### Historical precipitation, soil community legacies, and their impact on plant drought tolerance

Our hypothesis that historical precipitation patterns would impart legacy effects on soil microbiomes that influence plant growth under drought stress was supported. Bacterial and fungal compositional differences were apparent among the soils and inocula from prairie sites across a steep precipitation gradient in Kansas (Figures 1&2). We found that the interaction between the drought treatment and the mean annual precipitation of the soil inoculum source influenced shoot mass accumulation rate. This pattern held for alternative climate metrics (PDSI, aridity index, evapotranspiration, and soil moisture).

Following the stress gradient hypothesis (Bertness and Callaway 1994), we expected drought-stressed maize seedlings to perform better in the presence of microorganisms from historically dry soils, where chronic drought stress would have fostered positive interactions with neighboring plants. However, we found that plants performed better when the historic conditions of inocula were opposite from the contemporary conditions. The reason for this result is not immediately clear, but there are several possible explanations. Contemporary well-watered conditions may have relaxed the fitness consequences of expression of microbial traits that are critical under drier environments. For instance, microorganisms counteract the physiologic stress of drought by accumulating solutes, which reduces their internal water potential as a mechanism to avoid desiccation (Harris 1981). The accumulation of solutes requires considerable energy expenditure and constitutes a large sink of C and N resources (Killham and Firestone 1984; Schimel et al. 1989, 2007). The contemporary well-watered conditions relax these costly solute needs, perhaps allowing resources to be reallocated to functions that are beneficial to plants. Another possibility is that soils from historically wetter sites generally possess greater microbial biomass and diversity (Neilson et al. 2017; Araya et al. 2020; Wan et al. 2021), which is predicted to promote positive interactions and improved community function (Hooper et al. 2005). We found this to be the case for fungi, where our wettest prairie site showed the highest diversity, but this was less consistent for bacterial communities (Figure S3-4). A third potential explanation involves the balance of beneficial and pathogenic microbes in the soil community. Pathogen richness in soil, root, and shoot microbiomes is often positively associated with mean annual temperature and precipitation (Větrovský et al. 2019; Delgado-Baquerizo et al. 2020; Chen et al. 2021). Thus, soil communities from historically dry sites may have relatively fewer plant pathogens, resulting in greater plant performance under contemporary well-watered conditions. We cannot distinguish between these possible explanations using the data in hand; however, a future experiment that leverages rain exclusion structures to manipulate precipitation for a given site could effectively monitor changes in diversity and composition over time. In addition, this experimental manipulation strategy paired with a technique to quantify microbial biomass (e.g. quantitative stable-isotope probing) (Greenlon et al. 2022) could enable the exploration of the balance between beneficial, neutral, and pathogenic microbes.

### Microbial taxa associated with plant growth

Multiple bacteria and fungi had abundances correlated with plant responses to drought, with more taxa associated with shoot growth than root growth. This is likely the result of space limitation in the pots, as root mass accumulation rates tended to decrease across the experiment regardless of the drought treatment (Figure S19). Even so, we identified two families that were associated with both shoot and root mass, Acetobacteraceae and Corynebacteriaceae. Acetobacteraceae was associated with suppressed shoot growth but increased root growth. Although in our dataset Acetobacteraceae was represented mainly by the poorly-studied genus *Rhodopila*, this family contains many known diazotrophs, which may improve mineral nutrition of the plant (Reis and Teixeira 2015). Members of Acetobacteraceae, including the particularly well-studied *Gluconacetobacter diazotrophicus*, have been observed to colonize internal root tissues under both natural conditions and inoculation experiments (Luna et al. 2010; Rouws et al. 2010). This root colonization proficiency may help to explain the increased root mass we observed from this association; however, given the decreased shoot mass we also observed, Acetobacteraceae may have compartment-specific dynamics that warrant further investigation. In contrast, Corynebacteriaceae was associated with suppressed growth of both roots and shoots. Members of this family have been identified as root endophytes of multiple plant species (Stafford et al. 2005; Vik et al. 2013), with some showing phytopathogenic properties and specificity to maize (Vidaver and Mandel 1974; Dye and Kemp 1977). Additionally, during a multi-generation experimental evolution study, Corynebacteriaceae (among other bacterial families) was consistently associated with delayed flowering in multiple plant species (Panke-Buisse et al. 2015). Streptococcaceae was the only family that showed an association with plant growth that was modulated by the drought treatment, with this family’s relative abundance negatively correlated with shoot mass accumulation rate in well-watered plants but not in droughted plants. Less is known about the relevance of Streptococcaceae to plant growth. In *Zingiber officinale*, this family was significantly enriched in rhizomes with advanced stages of soft rot (Huang et al. 2022). However, Streptococcaceae is prevalent in soil and plants (Chuah et al. 2016; Wang et al. 2021), typically at higher abundance in aerial plant compartments than in the rhizosphere (Minervini et al. 2015; Yu et al. 2020). These families represent targets for direct manipulation and *in-planta* testing to better understand the mechanism responsible for the growth suppression we observed under well-watered conditions.

## Conclusions

Water is a critical and limiting resource for plant growth as well as a beneficial microbiome. As maize is cultivated widely across the United States, production will have to contend with more frequent episodes of drought as the global climate warms. Engineering plant microbiomes to assist in plant resilience under drought remains an ambitious, but worthwhile goal. Our work illustrates the differential responses of root and shoot growth allocation to drought in two well-studied maize genotypes and shows that bacterial, but not fungal, microbiota undergo restructuring with drought, with decreases in the relative abundance of drought-intolerant taxa. In contrast to predictions based on the stress gradient hypothesis, soil microbiomes from perennially dry environments along a natural precipitation gradient did not enhance plant growth under drought. Instead, plants under drought tended to produce more aboveground biomass with soil microbiomes from historically wetter sites. The bacterial and fungal groups associated with biomass can serve to direct future cultivation and *in-planta* testing on maize growth.

## Acknowledgements

We thank members of the Wagner lab for helpful comments in preparing this article. In addition, we thank Zachary Harris and Nichole Ginnan for helpful discussions. This work was funded by the United States Department of Agriculture National Institute of Food and Agriculture (USDA-NIFA) grant #2022-67013-36672 to MK and MRW, National Science Foundation grant #IOS-2016351 to MRW, Novo Nordisk Foundation INTERACT project under award number NNF19SA0059360 to MK, and seed funding from the NSF under award No. OIA-1656006 to Dr. Kristin Bowman-James and matching support from the State of Kansas through the Kansas Board of Regents. We thank the University of Kansas Medical Center – Genomics Core for generating the sequence data sets. The Genomics Core is supported by the Kansas Intellectual and Developmental Disabilities Research Center (NIH U54 HD 090216), the Molecular Regulation of Cell Development and Differentiation – COBRE (P30 GM122731), The NIH S10 High End Instrumentation Grant (NIH S10OD021743) and the Frontiers CTSA Grant (UL1TR002366).

## Data Availability

Raw sequencing data is available on NCBI SRA under BioProject ID PRJNA913622 and SRA accessions SAMN32302538-SAMN32303072 for root-associated samples and SAMN33863891-SAMN33864082 for soil and inoculum samples. Code to reproduce the analysis and figures are available on GitHub at https://github.com/Kenizzer/Maize_Drought_Microbiome.

## Author Contributions

BAS and PMH collected soil samples across Kansas. MRK, MK, and MRW designed the experiment and collection methods. MRK performed sample collection and processing. AC and NEF performed DNA extractions and library preparations. NEF and JFS performed the data analysis. JFS wrote the first draft of the manuscript. All authors contributed to data interpretation, the writing and reviewing of the manuscript, and approved the final draft.

## Supplemental Figures

**Figure S1.**
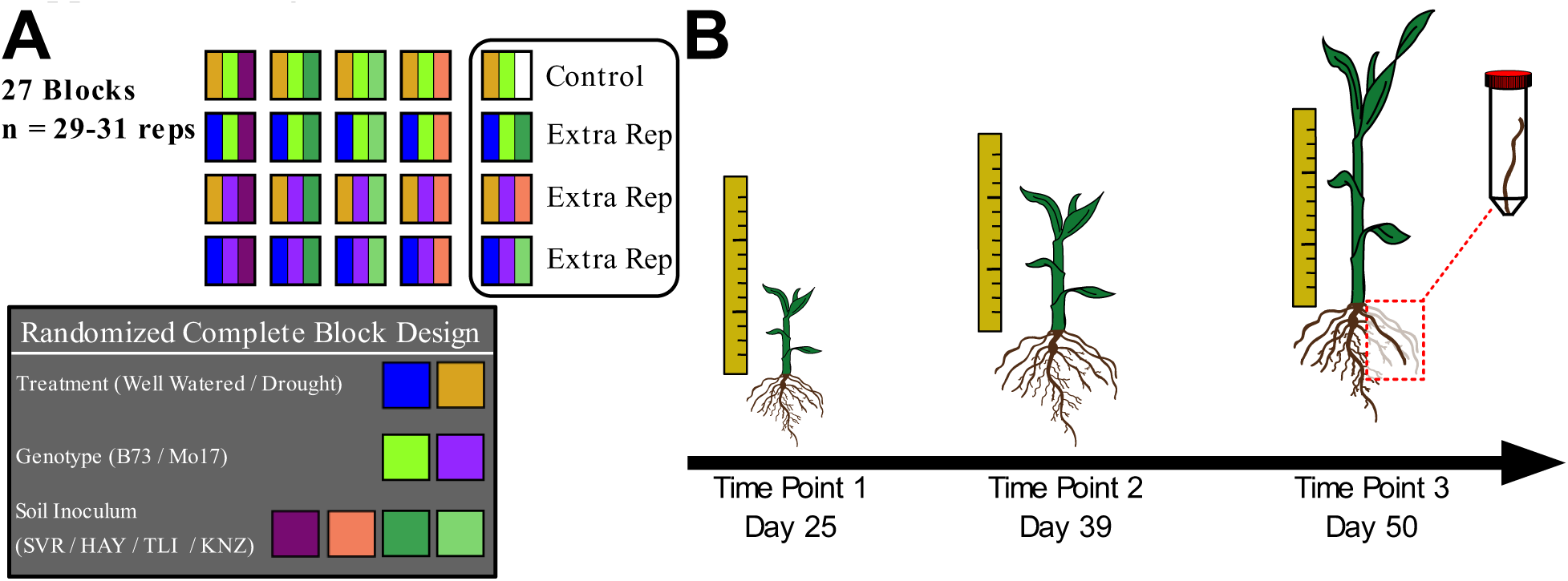
Experimental design. **A)** A complete randomized block design was used; depicted is a single representative block (number of blocks=27). Each pot was filled with sterile calcined clay soil mix and received a seed of either *Zea mays* genotype B73 or Mo17, was subject to either well-watered or drought conditions, and was inoculated with one of the four inocula. One uninoculated control (denoted with white stripe) and three additional randomized replicates were included per randomized block. **B)** Plant biomass was collected in a time series: for time point one (day 25) and two (day 39), 68 plants each were destructively harvested, then dried for root and shoot biomass. For the third time point (day 50), all remaining plants (n=379 – non-germinates) were harvested. Prior to drying for biomass measures, one nodal root from each plant was collected for amplicon sequencing for bacterial and fungal microbiome composition.

**Figure S2.**
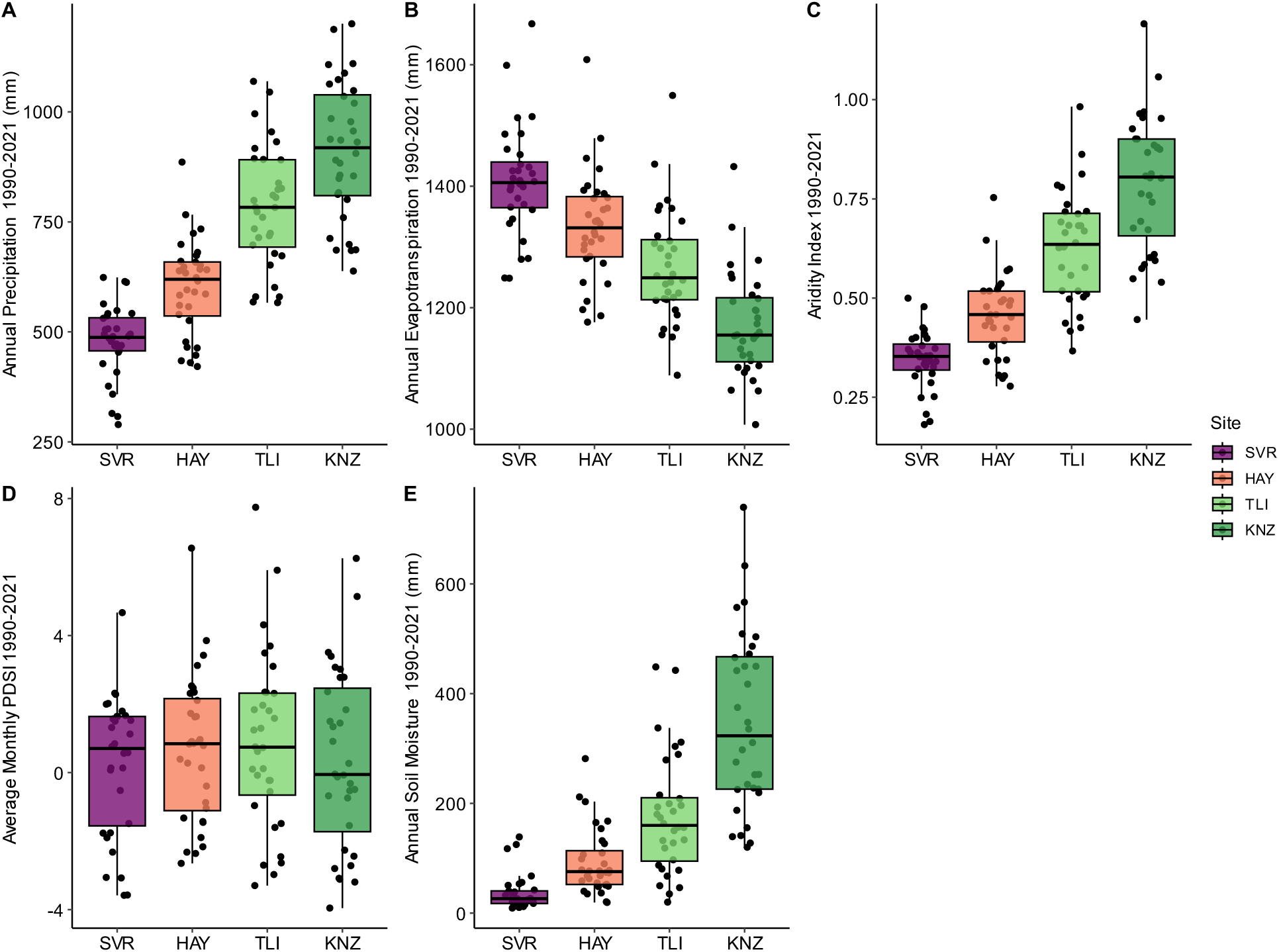
Climatic metrics for each collection site from 1990-2021. **A)** Annual precipitation, **B)** annual evapotranspiration, **C)** aridity index, **D)** average monthly Palmers Drought Stress Index (PDSI), and **E)** annual soil moisture. Boxplot hinges represent the 1^st^ and 3^rd^ quartiles; whiskers represent 1.5 times the interquartile range.

**Figure S3.**
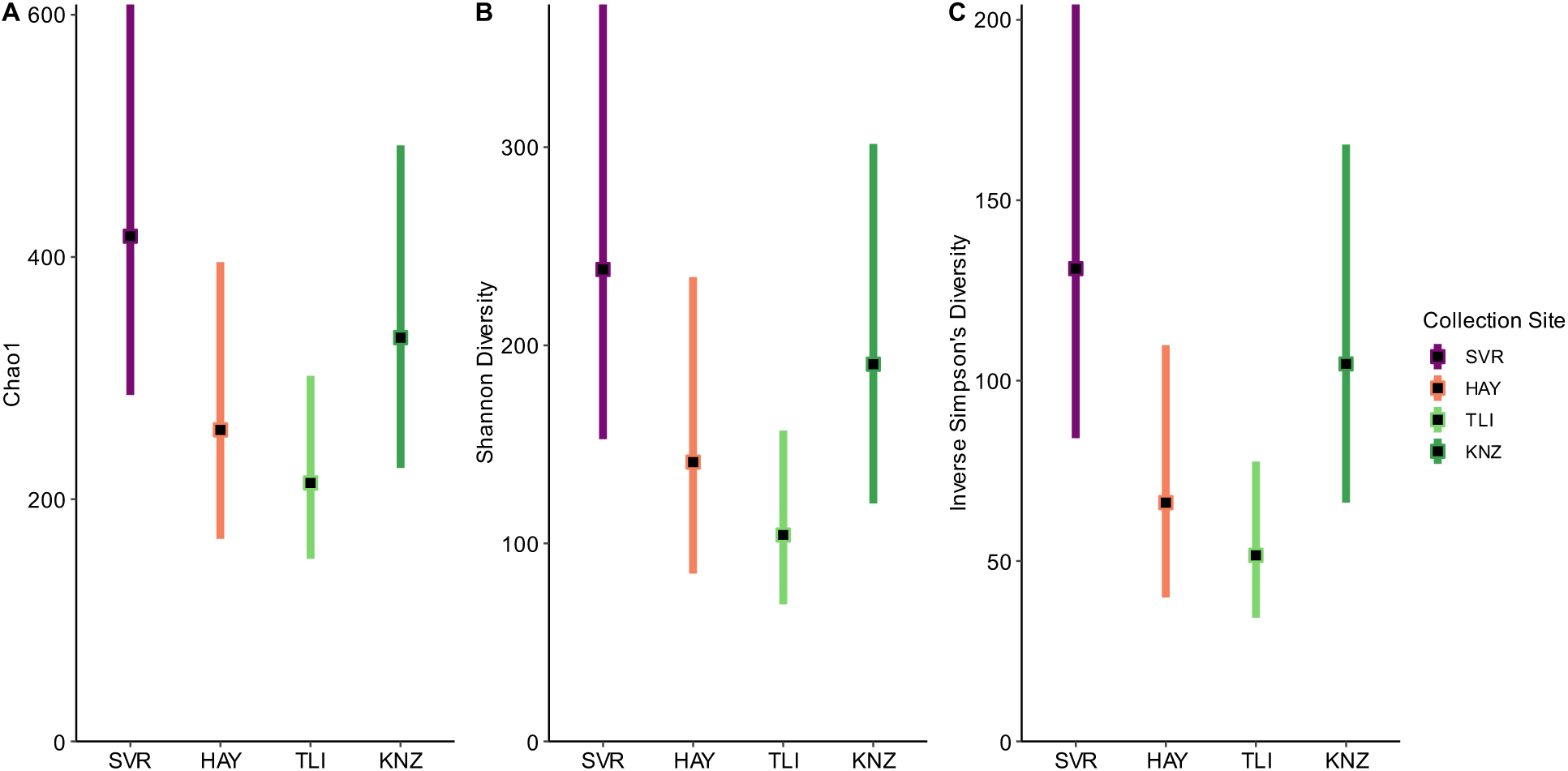
Bacterial alpha diversity values for **A)** Chao1 **B)** Shannon Diversity and **C)** Inverse Simpson’s Diversity. Black squares represent estimated marginal means (EMM) for each collection site and colored bars show the 95% confidence intervals for the EMMs.

**Figure S4.**
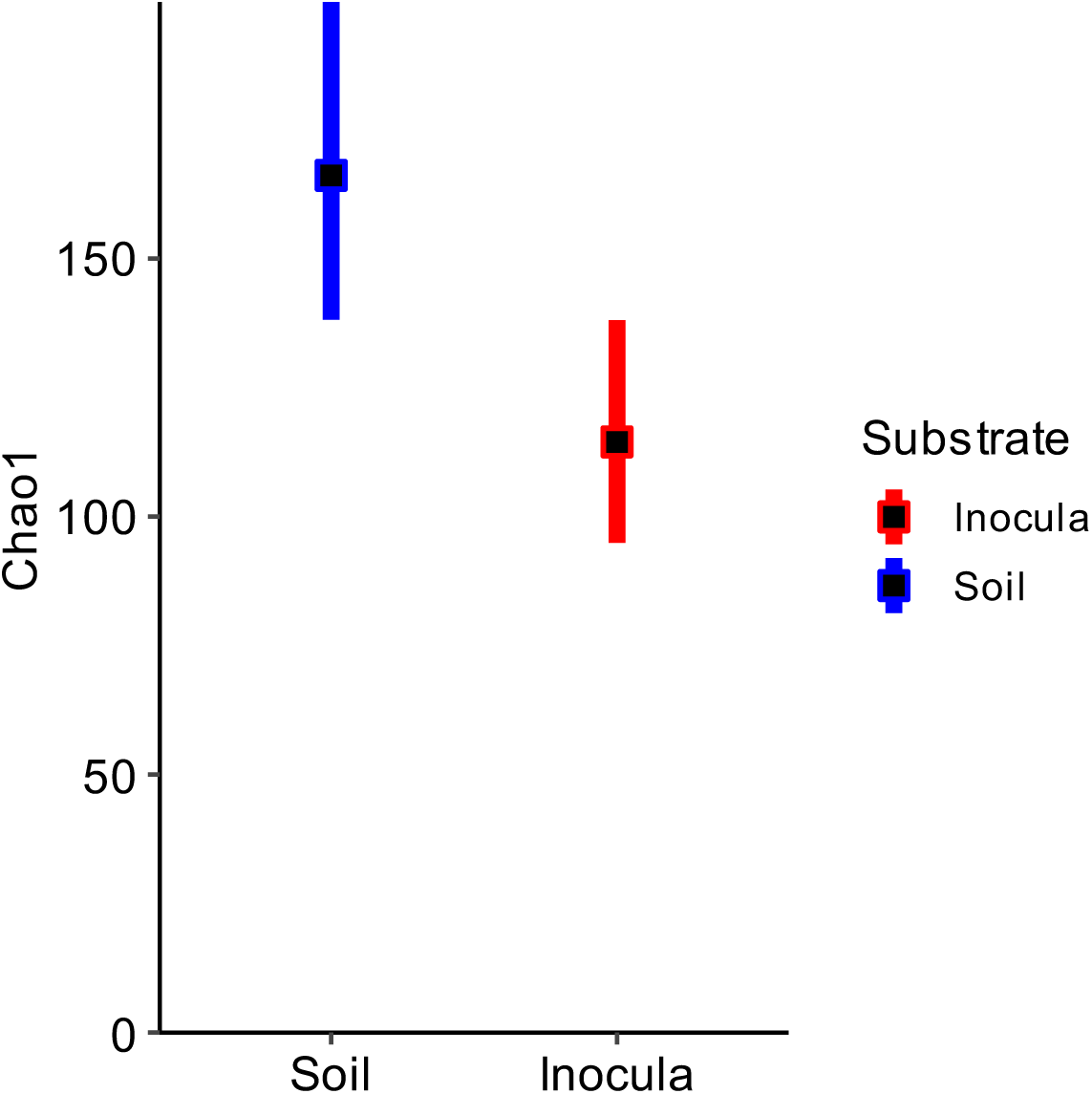
Fungal alpha diversity values for **A/C)** Chao1 and **B/D)** Inverse Simpson’s Diversity. Black squares represent estimated marginal means (EMM) for each collection site or substrate and colored bars show the 95% confidence intervals for the EMMs.

**Figure S5.**
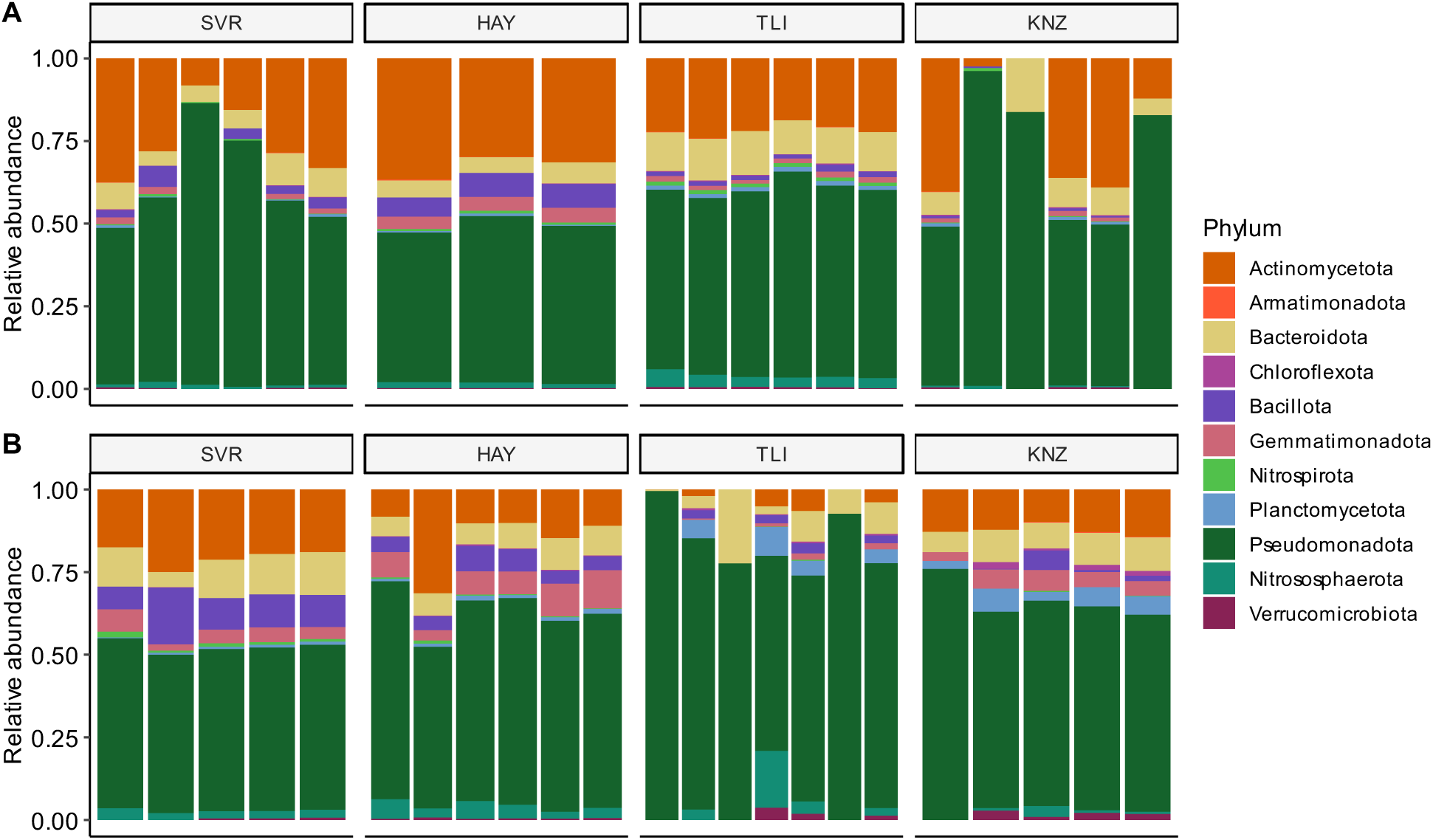
Taxonomic bar plots for bacterial phyla for **A)** soil and **B)** inocula samples. Historical annual precipitation for each collection site (SVR, HAY, TLI, KNZ) increases from left to right. Each sample is represented by a single line with the relative abundance (non CLR transformed data) of each phylum displayed by color.

**Figure S6.**
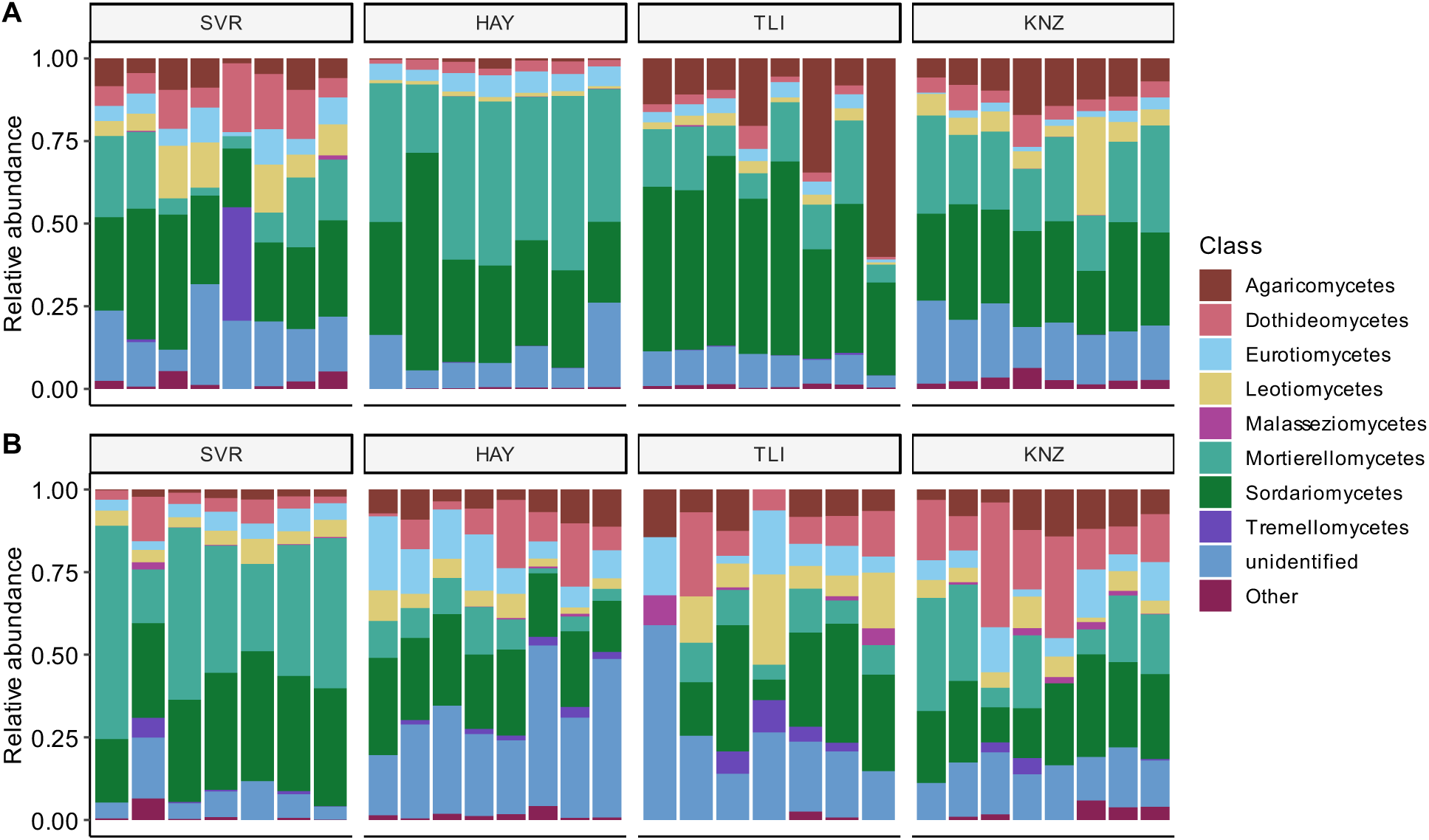
Taxonomic bar plots for fungal classes for **A)** soil and **B)** inocula samples. Historical annual precipitation for each collection site (SVR, HAY, TLI, KNZ) increases from left to right. Each sample is represented by a single line with the relative abundance (non CLR transformed data) of each class displayed by color.

**Figure S7.**
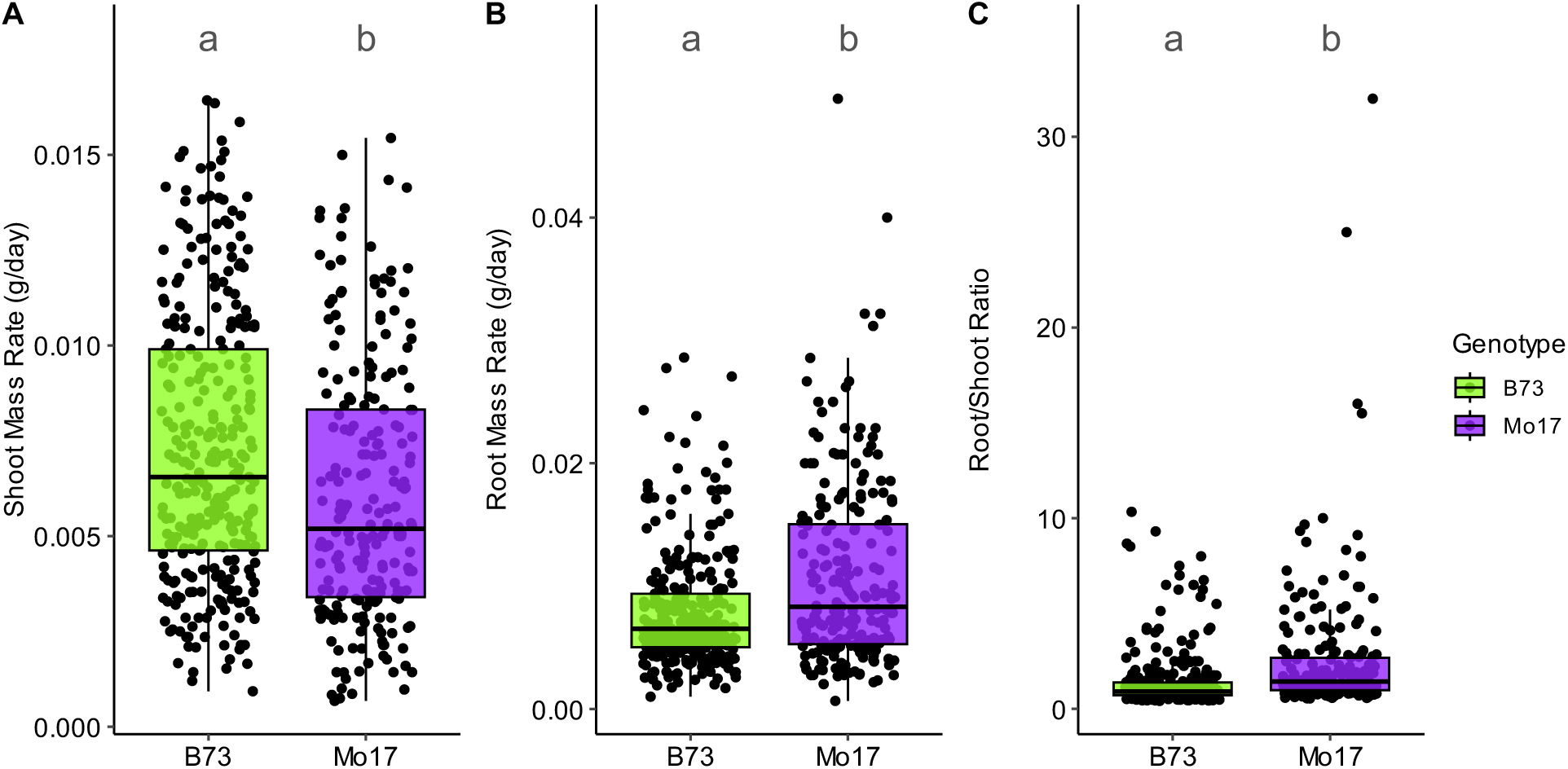
Maize genotypes growth differs across **A)** shoot mass rate, **B)** root mass rate, and **C)** root/shoot ratio. Boxplot hinges represent the 1^st^ and 3^rd^ quartiles; whiskers represent 1.5 times the interquartile range; letters above box plots denote significant differences in estimated marginal means (*p*-value=<0.01; see Table 1 for full ANOVA table).

**Figure S8.**
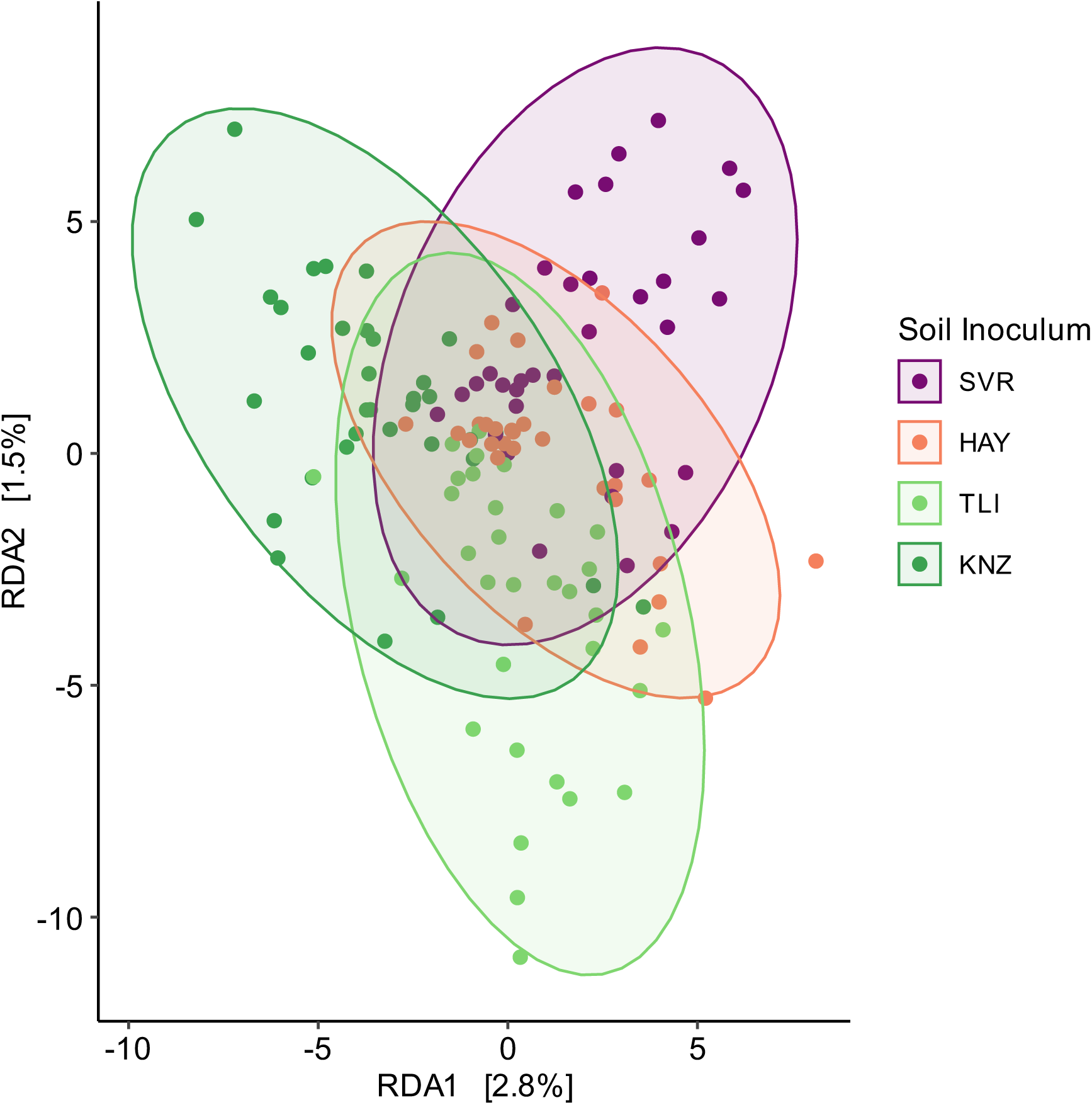
Redundancy analysis for root bacterial community composition. Points are colored by soil inoculum. Normal data ellipses are colored by soil inoculum assuming a multivariate normal distribution.

**Figure S9.**
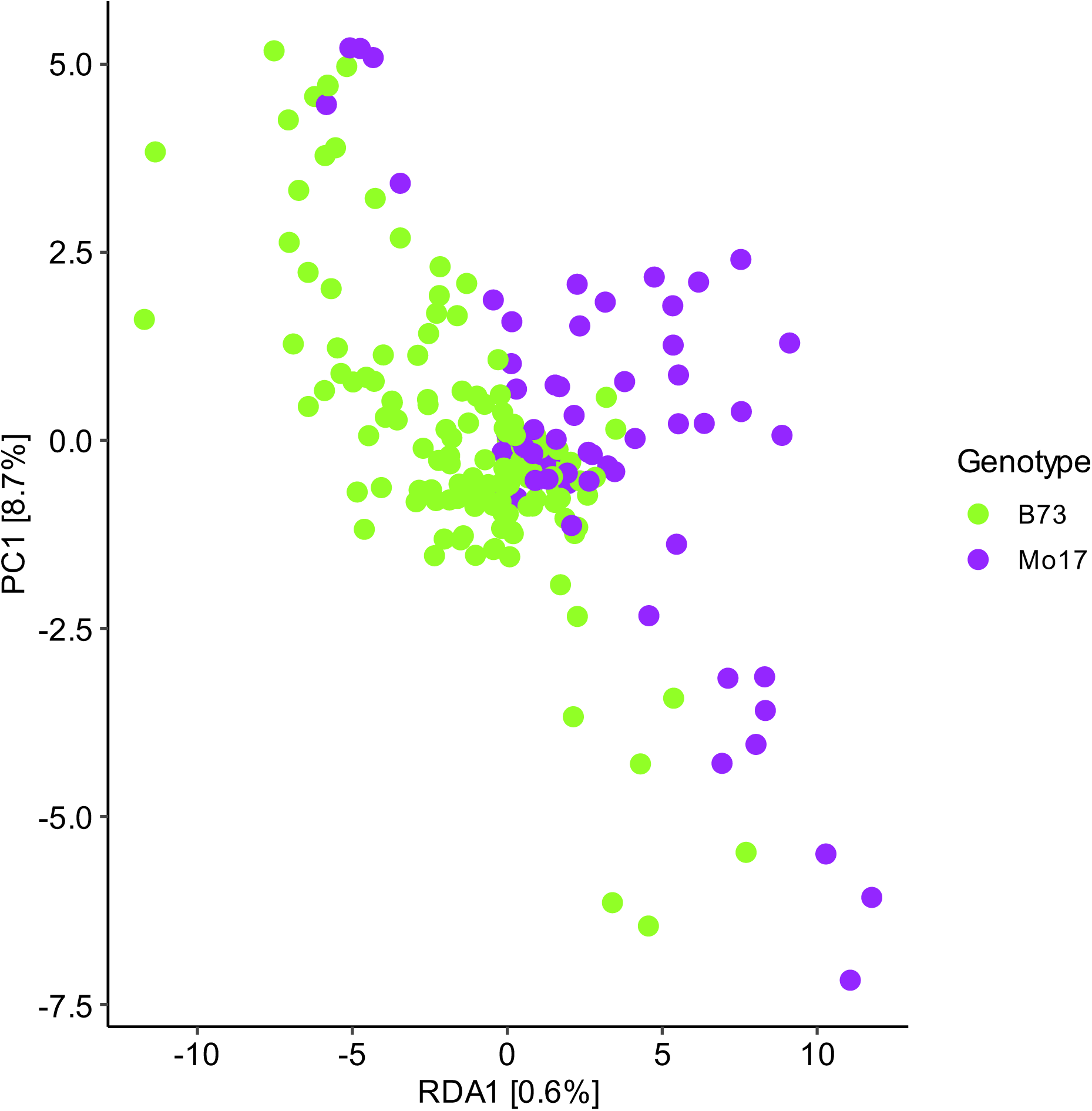
Redundancy analysis for root bacterial community composition. Points are colored by maize genotype.

**Figure S10.**
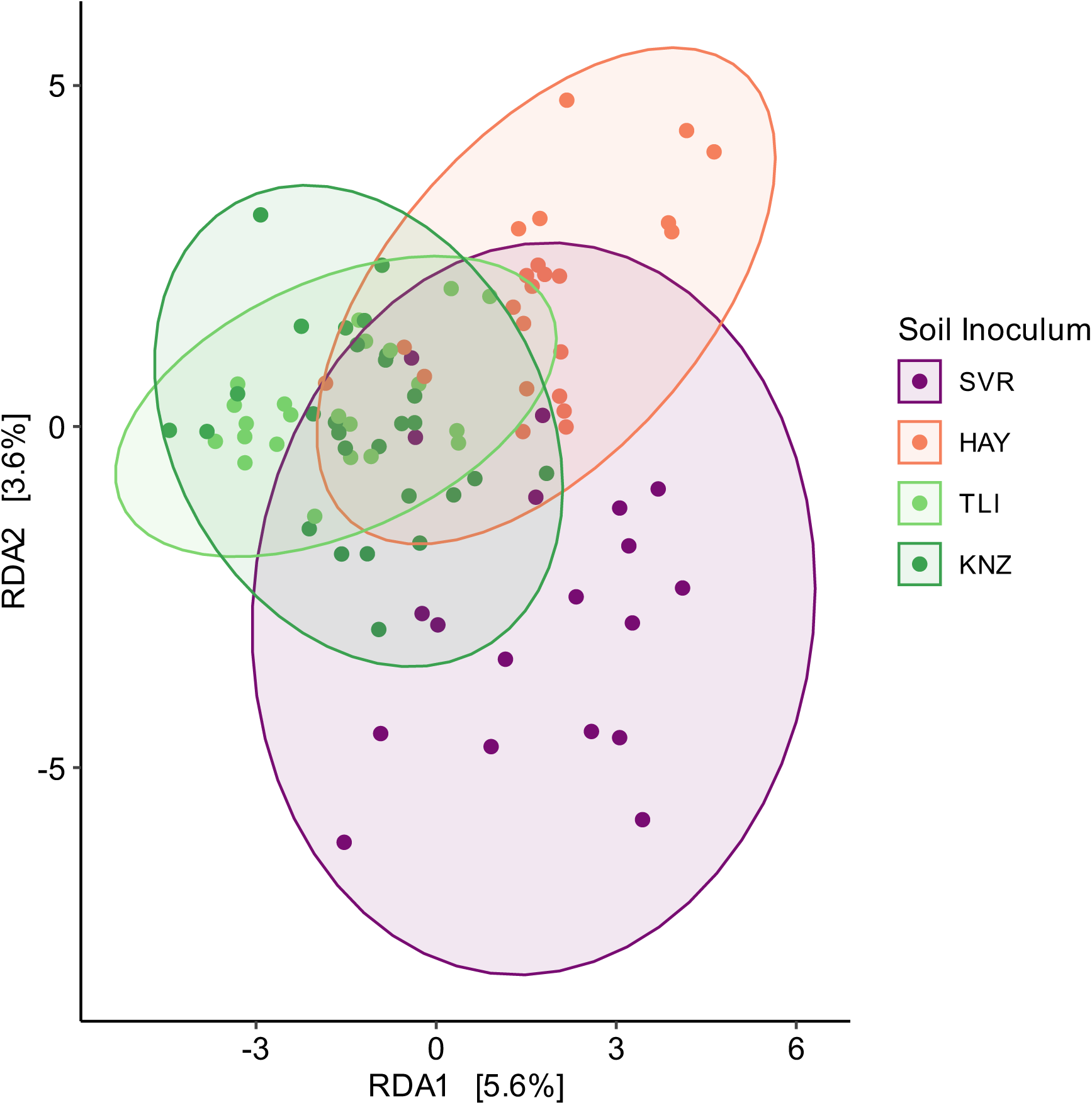
Redundancy analysis for root fungal community composition. Points are colored by soil inoculum. Normal data ellipses are colored by soil inoculum assuming a multivariate normal distribution.

**Figure S11.**
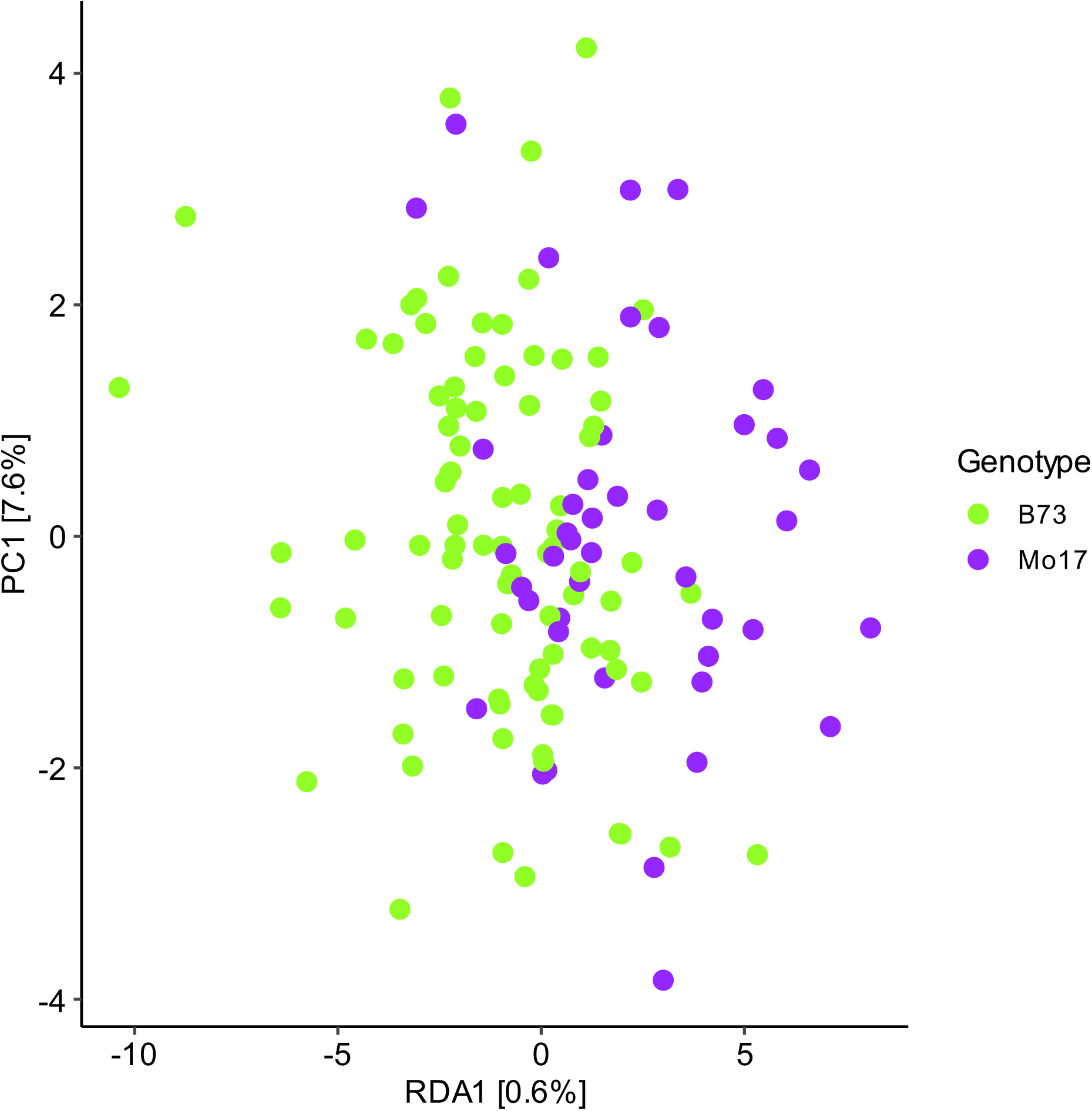
Redundancy analysis for fungal community composition. Points are colored by maize genotype.

**Figure S12.**
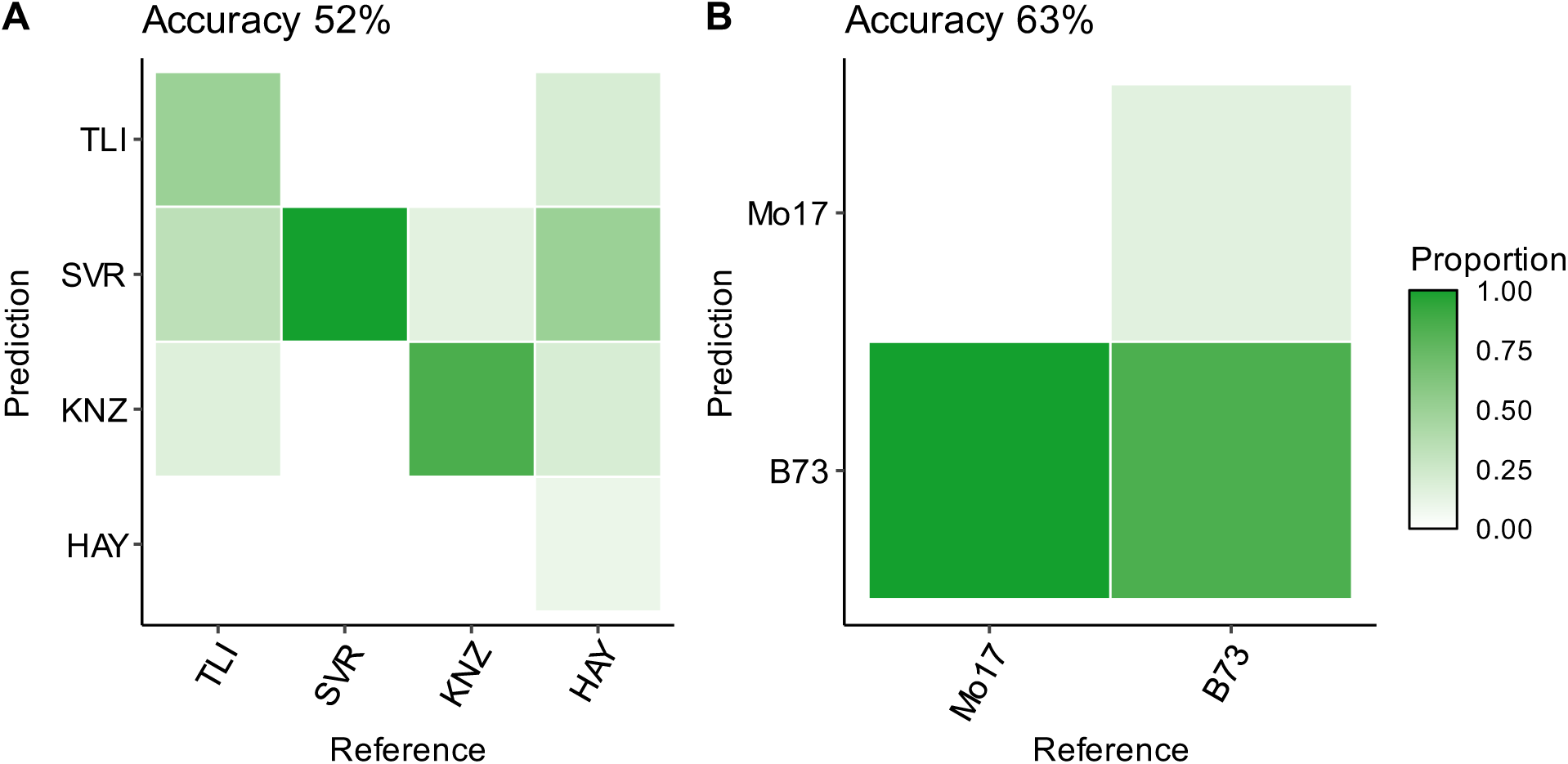
Confusion matrices depicting the classification accuracy of random forest classifiers trained on bacterial data predicting **A)** Soil Inoculum and **B)** Genotype. The models were trained using an 80:20 data split (80% train, 20% test) with 10-fold cross validation, shading represents the proportion of predictions for a label. Labels on the left represent the predicted labels and the labels on the bottom represent the actual labels. The model wide accuracy is given above each confusion matrix, class-wise statistics are provided in table in Table S8.

**Figure S13.**
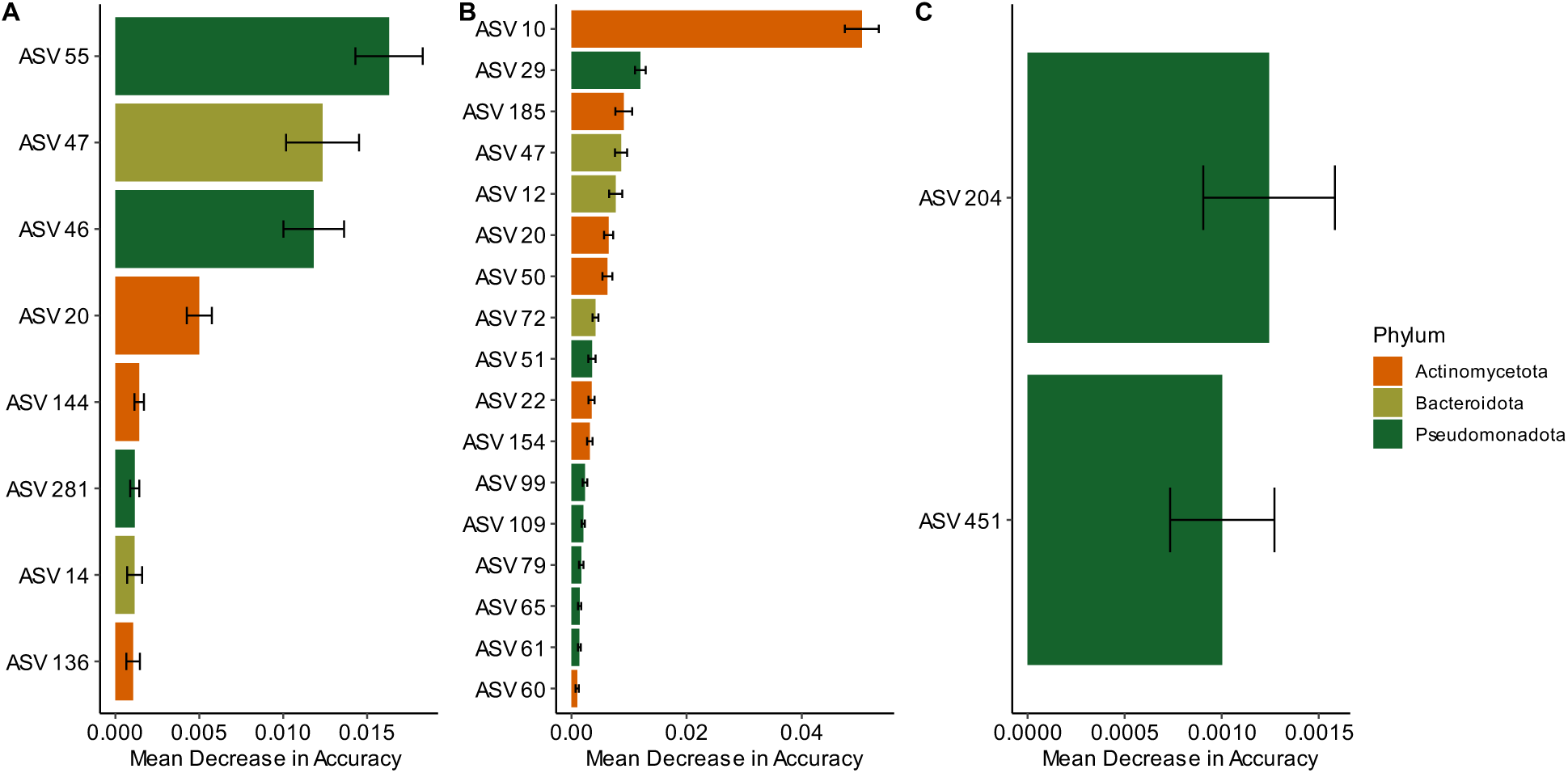
Bacterial ASV importance plots per random forest classifier. Results for **A)** Drought Treatment, **B)** Soil Inoculum, and **C)** Genotype classifiers. For each random forest classifier, 10-fold cross validation was used to assess the mean decrease in accuracy of the classifier when a particular ASV is withheld. Error bars represent the standard error across the 10-folds. Only ASVs with a mean decrease in accuracy greater than 0.001 are shown.

**Figure S14.**
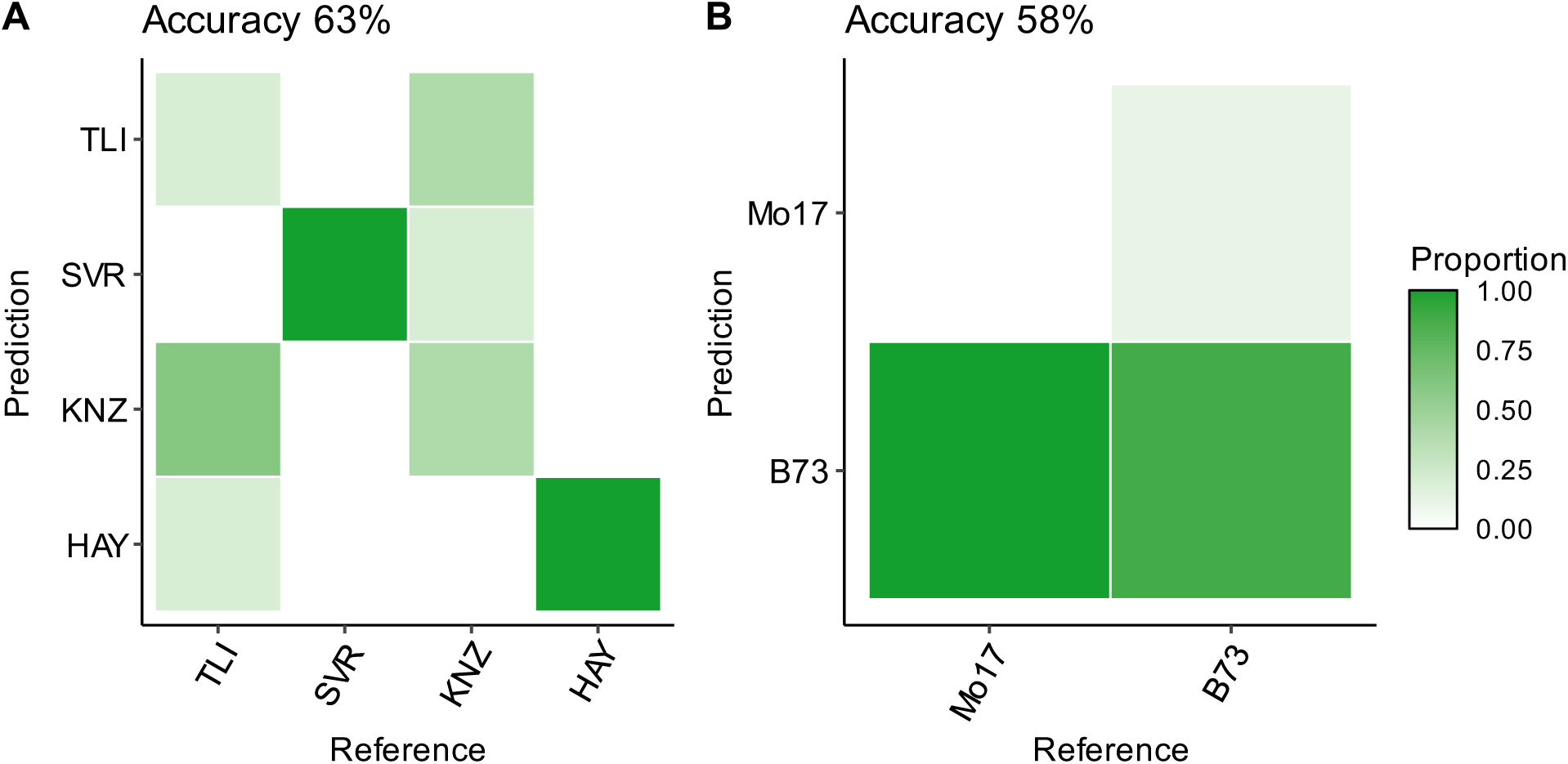
Confusion matrices depicting the classification accuracy of random forest classifiers trained on fungal data predicting **A)** Soil Inoculum and **B)** Genotype. The models were trained using an 80:20 data split (80% train, 20% test) with 10-fold cross validation, shading represents the proportion of predictions for a label. Labels on the left represent the predicted labels and the labels on the bottom represent the actual labels. The model wide accuracy is given above each confusion matrix, class-wise statistics are provided in table in Table S9.

**Figure S15.**
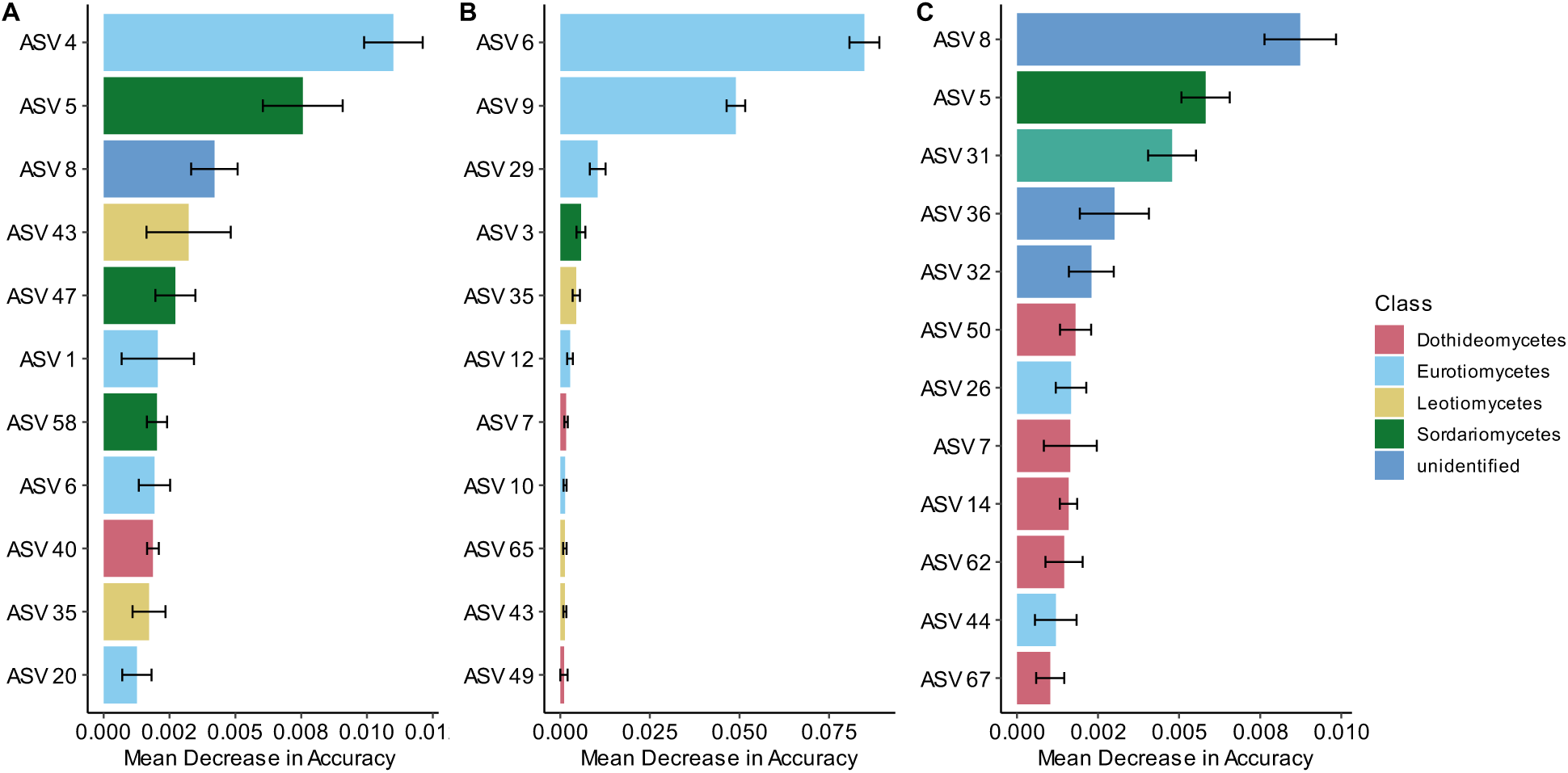
Fungal ASV importance plots per random forest classifier. Results for **A)** Drought Treatment, **B)** Soil Inoculum, and **C)** Genotype classifiers. For each random forest classifier, 10-fold cross validation was used to assess the mean decrease in accuracy of the classifier when a particular ASV is withheld. Error bars represent the standard error across the 10-folds. Only ASVs with a mean decrease in accuracy greater than 0.001 are shown.

**Figure S16.**
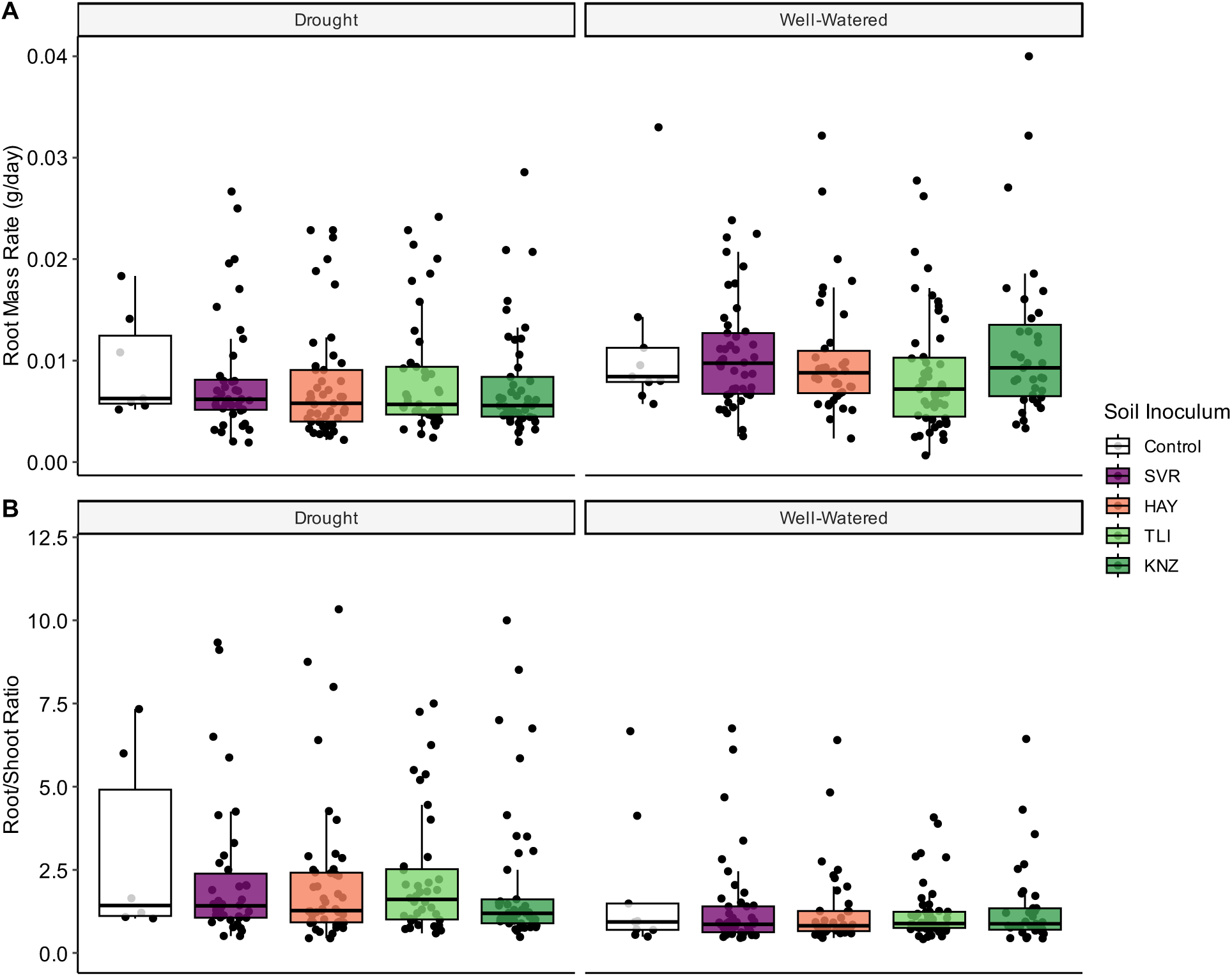
Root growth by soil inoculum. Boxplots are colored by soil inoculum for **A)** root mass rate and **B)** root/shoot ratio (three R/S data points were removed to reduce the y axis range and aid in visualization). Separate facets represent drought-stressed and well-watered plants. Boxplot hinges represent the 1^st^ and 3^rd^ quartiles; whiskers represent 1.5 times the interquartile range.

**Figure S17.**
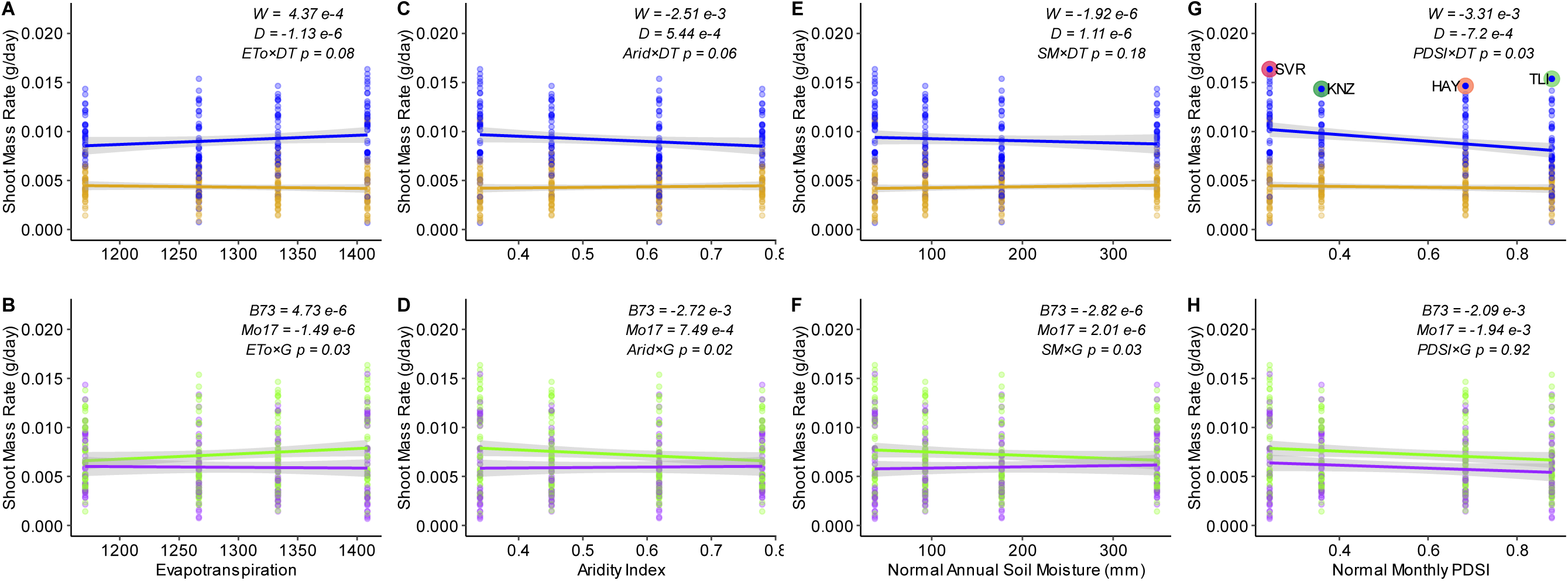
Shoot growth rate across climatic metrics. Trend lines for shoot mass accumulation rate against **A-B)** Evapotranspiration, **C-D)** Aridity Index, **E-F)** Soil Moisture, and **G-H)** Monthly Palmers Drought Stress Index (PDSI), for interactions with drought treatment and genotype, respectively, for plants inoculated with prairie soils. For PDSI, colors represent the prairie sites as PDSI values do not follow an east to west linear relationship (see Figure S2). For each regression line, grey shading represents the 95% confidence interval.

**Figure S18.**
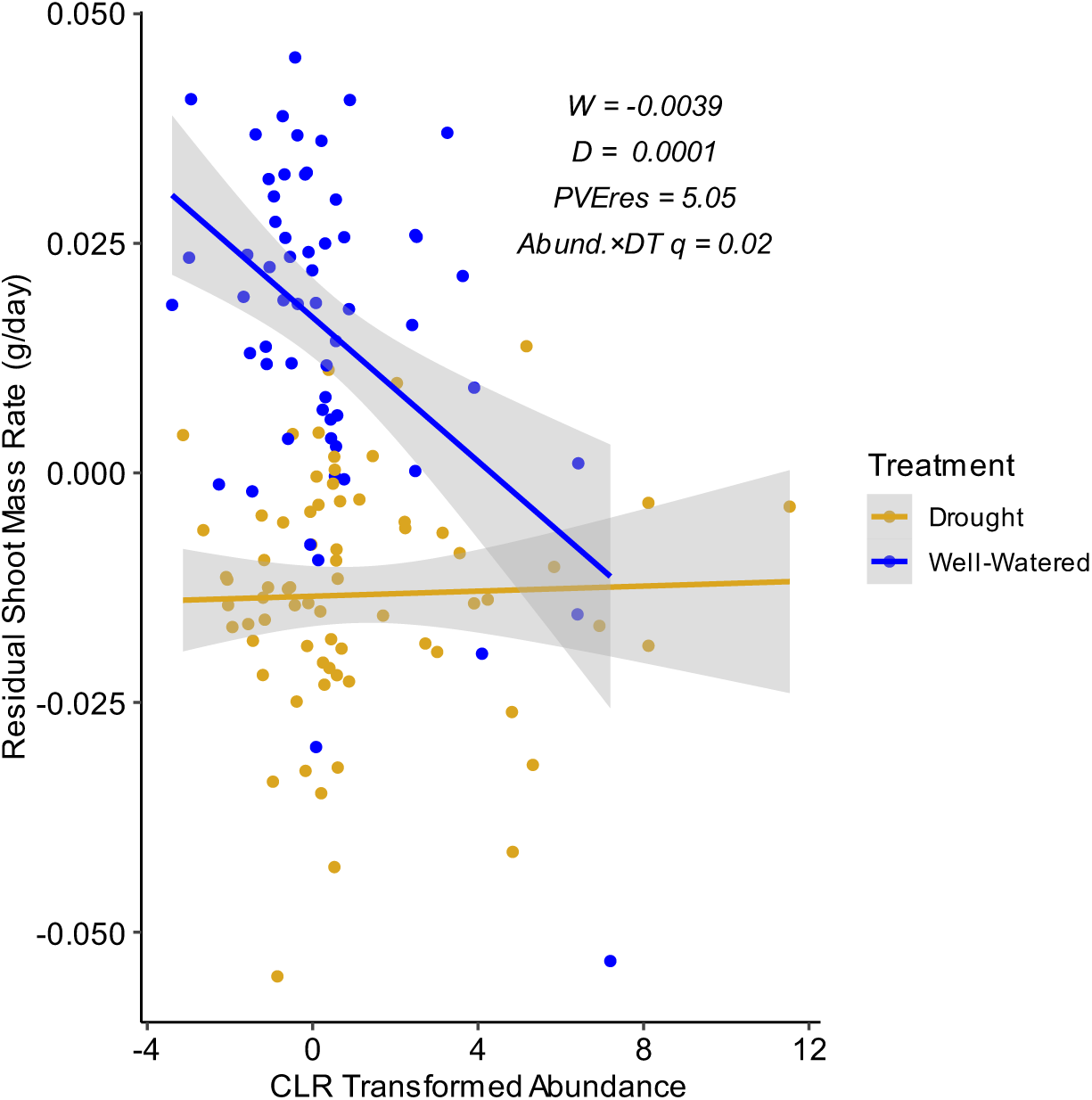
The bacterial family Streptococcaceae showed a significant interaction between CLR transformed abundance and drought treatment for residual shoot mass rate after accounting for other experimental and technical factors. For each regression line, grey shading represents the 95% confidence interval.

**Figure S19.**
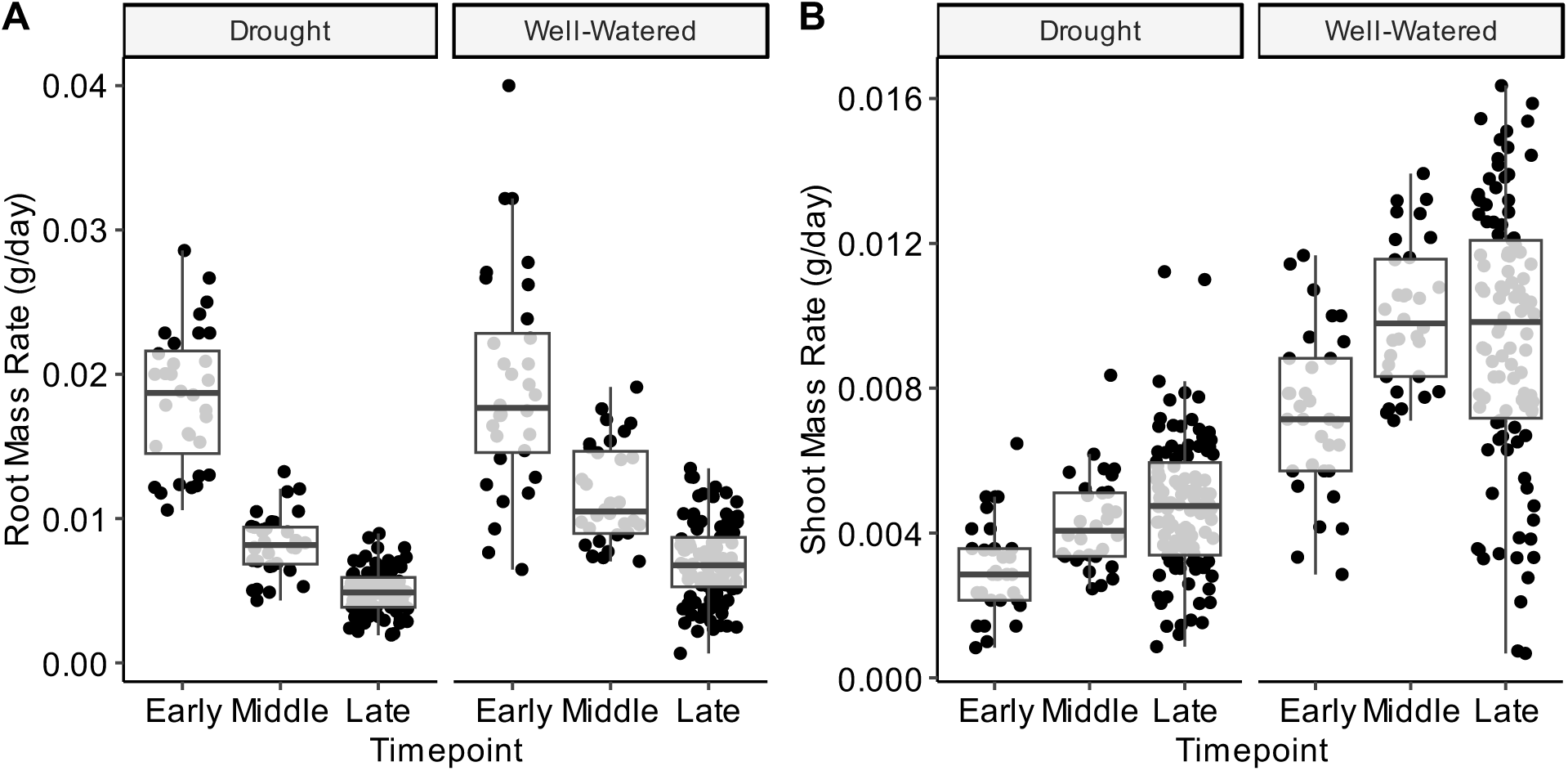
Plant growth rates across the collection time points of the study. **A**) Root mass rate tended to decrease over the course of the experiment and was modulated by the drought treatment (ANOVA; *‘Timepoint × Treatment’ p*-value = 0.02). **B**) Shoot mass rate tended to increase over the course of the experiment (ANOVA; *‘Timepoint’ p*-value< 0.01). Boxplot hinges represent the 1^st^ and 3^rd^ quartiles; whiskers represent 1.5 times the interquartile ran

## Supplemental Tables

**Table S1.** ANOVA table for bacterial alpha diversity metrics of soil samples collected from six locations across the Kansas precipitation gradient.

| Factor | DF | ln(Chao1) |  |  | ln(Shannon's Diversity) |  |  | ln(Inverse Simpson's Diversity) |  |  |
| --- | --- | --- | --- | --- | --- | --- | --- | --- | --- | --- |
|  |  | SS | F-value | p-value | SS | F-value | p-value | SS | F-value | p-value |
| Substrate | 1 | 1.12 | 2.94 | 0.09 | 2.17 | 4.09 | **0.05** | 2.01 | 3.83 | 0.06 |
| Collection Site | 3 | 8.36 | 7.32 | **<0.01** | 7.64 | 4.80 | **0.01** | 6.50 | 4.12 | **0.01** |
| Sequence Depth | 1 | 69.73 | 183.32 | **<0.01** | 50.99 | 96.14 | **<0.01** | 32.09 | 60.99 | **<0.01** |
| Residual | 38 | 14.46 |  |  | 20.15 |  |  | 19.99 |  |  |

Bold values denote significant effects, $p$ -value<0.05

**Table S2.** ANOVA table for fungal alpha diversity metrics of soil samples collected from six locations across the Kansas precipitation gradient.

| Factor | DF | Ln(Chao1) |  |  | ln(Shannon's Diversity) |  |  | ln(Inverse Simpson's Diversity) |  |  |
| --- | --- | --- | --- | --- | --- | --- | --- | --- | --- | --- |
|  |  | SS | F-value | p-value | SS | F-value | p-value | SS | F-value | p-value |
| Substrate | 1 | 8.58 | 36.27 | **<0.01** | 0.02 | 0.06 | 0.80 | 0.85 | 1.87 | 0.18 |
| Collection Site | 3 | 0.84 | 1.19 | 0.32 | 2.25 | 2.08 | 0.11 | 3.46 | 2.55 | 0.06 |
| Sequence Depth | 1 | 10.89 | 46.01 | **<0.01** | 0.03 | 0.08 | 0.78 | 2.41 | 5.32 | **0.02** |
| Residual | 55 | 13.01 |  |  | 19.79 |  |  | 24.88 |  |  |

Bold values denote significant effects, $p$ -value<0.05

**Table S3.**
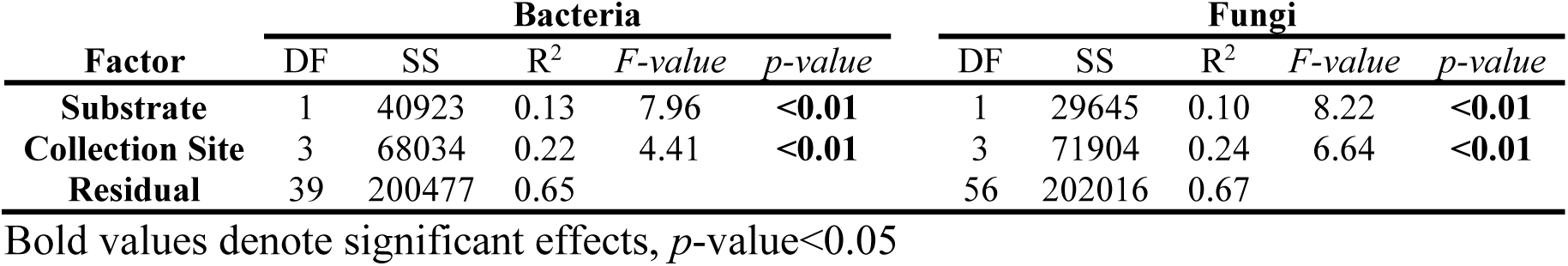
PERMANOVA table for bacterial and fungal community composition of soil and inocula samples collected from four locations across the Kansas precipitation gradient.

**Table S4.**
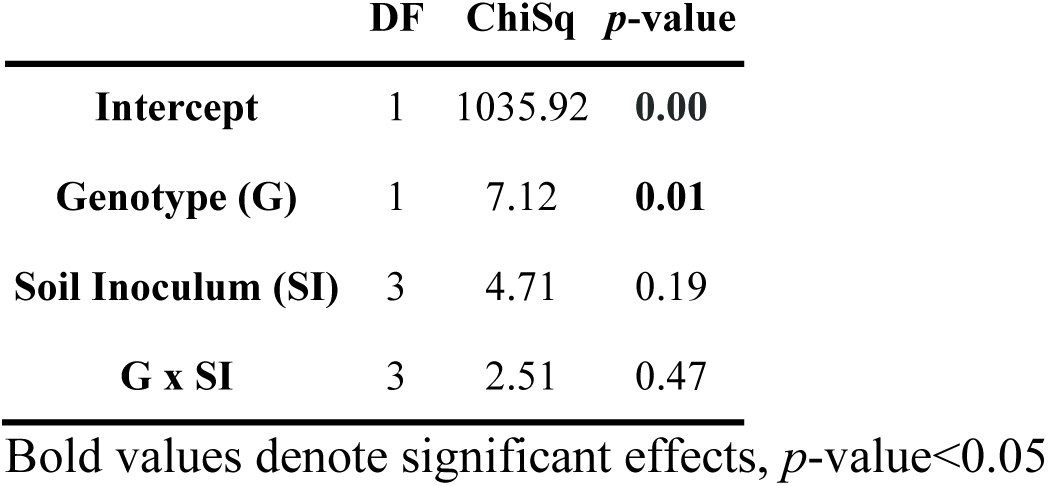
ANOVA table for maize emergence rate.

|  | DF | ChiSq | p-value |
| --- | --- | --- | --- |
| Intercept | 1 | 1035.92 | **0.00** |
| Genotype (G) | 1 | 7.12 | **0.01** |
| Soil Inoculum (SI) | 3 | 4.71 | 0.19 |
| G x SI | 3 | 2.51 | 0.47 |

Bold values denote significant effects, $p$ -value<0.05

**Table S5.** ANOVA table for bacterial alpha diversity metrics of maize nodal roots.

| Factor | DF | Chao1 |  |  | ln(Shannon's Diversity) |  |  | Ln(Inverse Simpson's Diversity) |  |  |
| --- | --- | --- | --- | --- | --- | --- | --- | --- | --- | --- |
|  |  | SS | F-value | p-value | SS | F-value | p-value | SS | F-value | p-value |
| Genotype (G) | 1 | 11476 | 0.60 | 0.44 | 0.06 | 0.13 | 0.72 | 0.04 | 0.08 | 0.77 |
| Soil Inoculum (SI) | 3 | 179893 | 3.11 | **0.03** | 0.88 | 0.61 | 0.61 | 0.09 | 0.07 | 0.98 |
| Drought Treatment (DT) | 1 | 55466 | 2.88 | 0.09 | 0.20 | 0.42 | 0.52 | 0.62 | 1.42 | 0.24 |
| G X SI | 3 | 79499 | 1.37 | 0.25 | 0.88 | 0.61 | 0.61 | 0.78 | 0.59 | 0.62 |
| G X DT | 1 | 25967 | 1.35 | 0.25 | 0.28 | 0.57 | 0.45 | 0.24 | 0.55 | 0.46 |
| SI x DT | 3 | 34773 | 0.60 | 0.62 | 0.66 | 0.46 | 0.71 | 0.43 | 0.33 | 0.80 |
| G X SI X DT | 3 | 6233 | 0.11 | 0.96 | 0.07 | 0.05 | 0.99 | 0.23 | 0.18 | 0.91 |

Bold values denote significant effects, $p$ -value<0.05

**Table S6.** ANOVA table for fungal alpha diversity metrics of maize nodal roots.

| Factor | DF | Chao1 |  |  | ln(Shannon's Diversity) |  |  | ln(Inverse Simpson's Diversity) |  |  |
| --- | --- | --- | --- | --- | --- | --- | --- | --- | --- | --- |
|  |  | SS | F-value | p-value | SS | F-value | p-value | SS | F-value | p-value |
| Genotype (G) | 1 | 107.32 | 0.59 | 0.45 | 0.56 | 1.62 | 0.21 | 0.71 | 2.47 | 0.12 |
| Soil Inoculum (SI) | 3 | 428.46 | 0.78 | 0.51 | 0.30 | 0.29 | 0.83 | 0.17 | 0.20 | 0.90 |
| Drought Treatment (DT) | 1 | 10.54 | 0.06 | 0.81 | 1.73 | 5.00 | **0.03** | 1.79 | 6.17 | **0.02** |
| G X SI | 3 | 805.97 | 1.47 | 0.23 | 0.53 | 0.51 | 0.68 | 0.63 | 0.72 | 0.54 |
| G X DT | 1 | 41.51 | 0.23 | 0.64 | 0.20 | 0.57 | 0.45 | 0.37 | 1.27 | 0.26 |
| SI x DT | 3 | 1044.15 | 1.90 | 0.14 | 2.71 | 2.60 | 0.06 | 2.12 | 2.44 | 0.07 |
| G X SI X DT | 3 | 557.96 | 1.01 | 0.39 | 1.05 | 1.01 | 0.39 | 1.06 | 1.22 | 0.31 |
| Genotype (G) | 1 | 107.32 | 0.59 | 0.45 | 0.56 | 1.62 | 0.21 | 0.71 | 2.47 | 0.12 |

Bold values denote significant effects, $p$ -value<0.05

**Table S7.** Optimal hyperparameters for bacterial and fungal machine learning classifiers trained to predict treatment, soil inoculum, and genotype.

| Factor | Bacteria |  | Fungi |  |
| --- | --- | --- | --- | --- |
|  | mtry | Minimum Node Side | mtry | Minimum Node Side |
| Treatment | 559 | 10 | 45 | 10 |
| Soil Inoculum | 437 | 5 | 66 | 1 |
| Genotype | 559 | 5 | 10 | 1 |

**Table S8.** Output statistics for bacterial machine learning models predicting treatment, genotype, and soil inoculum.

| Metric | Treatment | Genotype | Soil Inoculum |  |  |  |
| --- | --- | --- | --- | --- | --- | --- |
|  |  |  | SVR | HAY | TLI | KNZ |
| Precision | 0.700 | 0.708 | 0.333 | 1.000 | 0.600 | 0.667 |
| Recall | 0.875 | 0.850 | 1.000 | 0.100 | 0.500 | 0.857 |
| F1 | 0.778 | 0.773 | 0.500 | 0.182 | 0.546 | 0.750 |
| Prevalence | 0.593 | 0.741 | 0.148 | 0.370 | 0.222 | 0.259 |
| Balanced Accuracy | 0.665 | 0.425 | 0.826 | 0.550 | 0.702 | 0.854 |

**Table S9.** Output statistics for fungal machine learning models predicting treatment, genotype, and soil inoculum.

| Metric | Treatment | Genotype | Soil Inoculum |  |  |  |
| --- | --- | --- | --- | --- | --- | --- |
|  |  |  | SVR | HAY | TLI | KNZ |
| Precision | 0.500 | 0.625 | 0.667 | 0.875 | 0.333 | 0.400 |
| Recall | 0.400 | 0.882 | 1.000 | 1.000 | 0.200 | 0.400 |
| F1 | 0.444 | 0.732 | 0.800 | 0.933 | 0.250 | 0.400 |
| Prevalence | 0.526 | 0.654 | 0.105 | 0.368 | 0.263 | 0.263 |
| Balanced Accuracy | 0.478 | 0.441 | 0.971 | 0.958 | 0.529 | 0.593 |

**Table S10.**
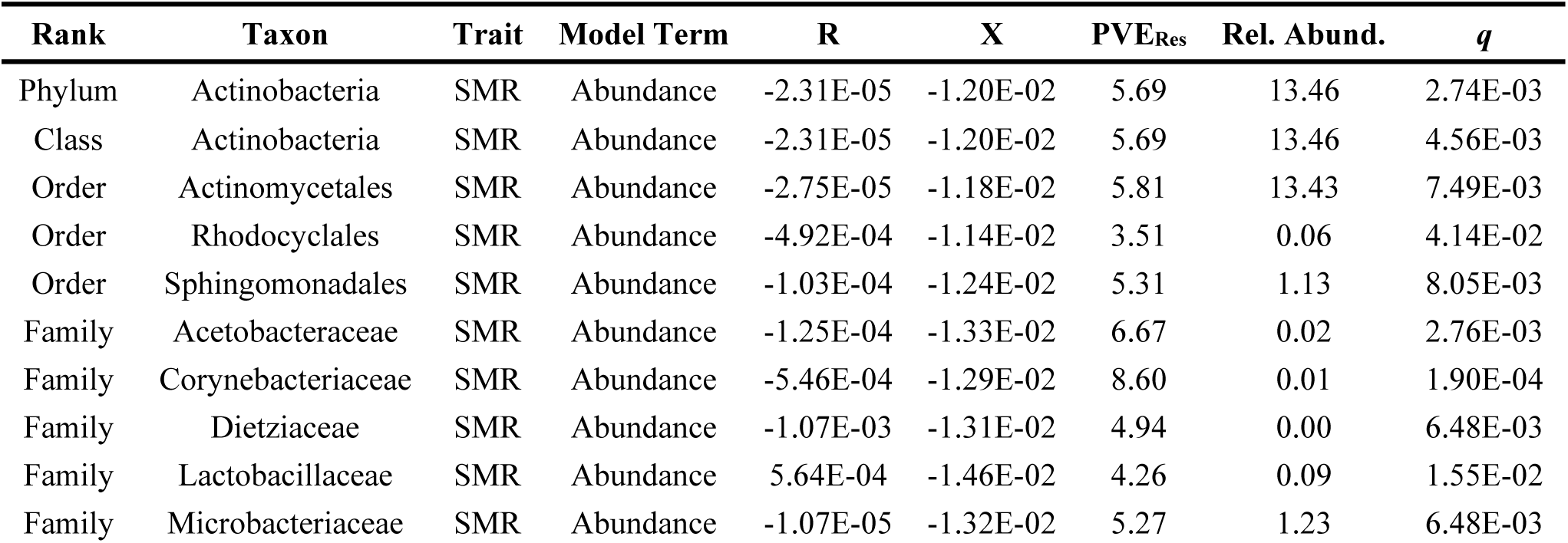
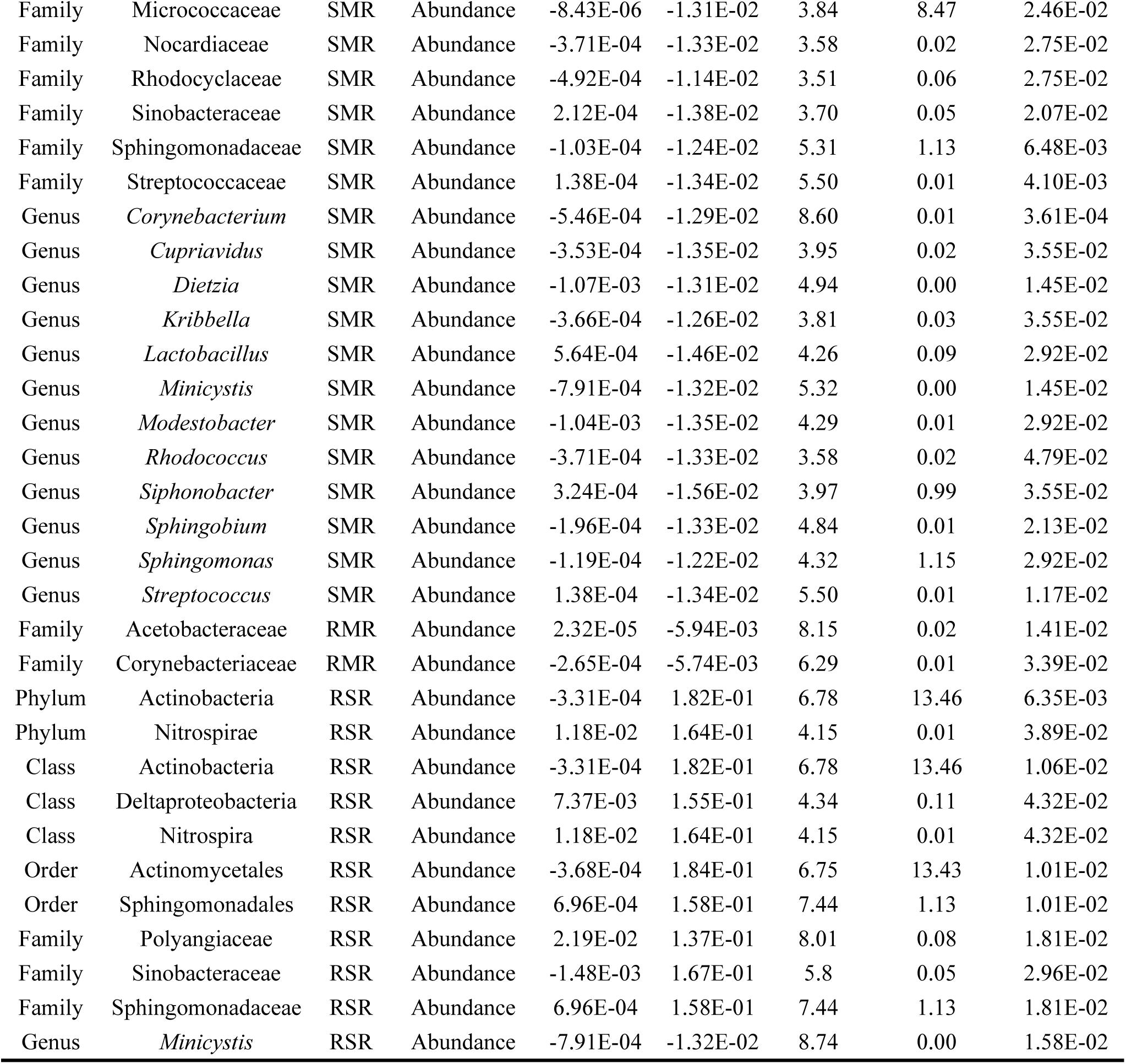
Bacterial taxa across taxonomic levels associated with square root transformed shoot mass rate.

| Rank | Taxon | Trait | Model Term | R | X | PVE<sub>Res</sub> | Rel. Abund. | q |
| --- | --- | --- | --- | --- | --- | --- | --- | --- |
| Phylum | Actinobacteria | SMR | Abundance | -2.31E-05 | -1.20E-02 | 5.69 | 13.46 | 2.74E-03 |
| Class | Actinobacteria | SMR | Abundance | -2.31E-05 | -1.20E-02 | 5.69 | 13.46 | 4.56E-03 |
| Order | Actinomycetales | SMR | Abundance | -2.75E-05 | -1.18E-02 | 5.81 | 13.43 | 7.49E-03 |
| Order | Rhodocyclales | SMR | Abundance | -4.92E-04 | -1.14E-02 | 3.51 | 0.06 | 4.14E-02 |
| Order | Sphingomonadales | SMR | Abundance | -1.03E-04 | -1.24E-02 | 5.31 | 1.13 | 8.05E-03 |
| Family | Acetobacteraceae | SMR | Abundance | -1.25E-04 | -1.33E-02 | 6.67 | 0.02 | 2.76E-03 |
| Family | Corynebacteriaceae | SMR | Abundance | -5.46E-04 | -1.29E-02 | 8.60 | 0.01 | 1.90E-04 |
| Family | Dietziaceae | SMR | Abundance | -1.07E-03 | -1.31E-02 | 4.94 | 0.00 | 6.48E-03 |
| Family | Lactobacillaceae | SMR | Abundance | 5.64E-04 | -1.46E-02 | 4.26 | 0.09 | 1.55E-02 |
| Family | Microbacteriaceae | SMR | Abundance | -1.07E-05 | -1.32E-02 | 5.27 | 1.23 | 6.48E-03 |
| Family | Micrococcaceae | SMR | Abundance | -8.43E-06 | -1.31E-02 | 3.84 | 8.47 | 2.46E-02 |
| Family | Nocardiaceae | SMR | Abundance | -3.71E-04 | -1.33E-02 | 3.58 | 0.02 | 2.75E-02 |
| Family | Rhodocyclaceae | SMR | Abundance | -4.92E-04 | -1.14E-02 | 3.51 | 0.06 | 2.75E-02 |
| Family | Sinobacteraceae | SMR | Abundance | 2.12E-04 | -1.38E-02 | 3.70 | 0.05 | 2.07E-02 |
| Family | Sphingomonadaceae | SMR | Abundance | -1.03E-04 | -1.24E-02 | 5.31 | 1.13 | 6.48E-03 |
| Family | Streptococcaceae | SMR | Abundance | 1.38E-04 | -1.34E-02 | 5.50 | 0.01 | 4.10E-03 |
| Genus | *Corynebacterium* | SMR | Abundance | -5.46E-04 | -1.29E-02 | 8.60 | 0.01 | 3.61E-04 |
| Genus | *Cupriavidus* | SMR | Abundance | -3.53E-04 | -1.35E-02 | 3.95 | 0.02 | 3.55E-02 |
| Genus | *Dietzia* | SMR | Abundance | -1.07E-03 | -1.31E-02 | 4.94 | 0.00 | 1.45E-02 |
| Genus | *Kribbella* | SMR | Abundance | -3.66E-04 | -1.26E-02 | 3.81 | 0.03 | 3.55E-02 |
| Genus | *Lactobacillus* | SMR | Abundance | 5.64E-04 | -1.46E-02 | 4.26 | 0.09 | 2.92E-02 |
| Genus | *Minicystis* | SMR | Abundance | -7.91E-04 | -1.32E-02 | 5.32 | 0.00 | 1.45E-02 |
| Genus | *Modestobacter* | SMR | Abundance | -1.04E-03 | -1.35E-02 | 4.29 | 0.01 | 2.92E-02 |
| Genus | *Rhodococcus* | SMR | Abundance | -3.71E-04 | -1.33E-02 | 3.58 | 0.02 | 4.79E-02 |
| Genus | *Siphonobacter* | SMR | Abundance | 3.24E-04 | -1.56E-02 | 3.97 | 0.99 | 3.55E-02 |
| Genus | *Sphingobium* | SMR | Abundance | -1.96E-04 | -1.33E-02 | 4.84 | 0.01 | 2.13E-02 |
| Genus | *Sphingomonas* | SMR | Abundance | -1.19E-04 | -1.22E-02 | 4.32 | 1.15 | 2.92E-02 |
| Genus | *Streptococcus* | SMR | Abundance | 1.38E-04 | -1.34E-02 | 5.50 | 0.01 | 1.17E-02 |
| Family | Acetobacteraceae | RMR | Abundance | 2.32E-05 | -5.94E-03 | 8.15 | 0.02 | 1.41E-02 |
| Family | Corynebacteriaceae | RMR | Abundance | -2.65E-04 | -5.74E-03 | 6.29 | 0.01 | 3.39E-02 |
| Phylum | Actinobacteria | RSR | Abundance | -3.31E-04 | 1.82E-01 | 6.78 | 13.46 | 6.35E-03 |
| Phylum | Nitrospirae | RSR | Abundance | 1.18E-02 | 1.64E-01 | 4.15 | 0.01 | 3.89E-02 |
| Class | Actinobacteria | RSR | Abundance | -3.31E-04 | 1.82E-01 | 6.78 | 13.46 | 1.06E-02 |
| Class | Deltaproteobacteria | RSR | Abundance | 7.37E-03 | 1.55E-01 | 4.34 | 0.11 | 4.32E-02 |
| Class | Nitrospira | RSR | Abundance | 1.18E-02 | 1.64E-01 | 4.15 | 0.01 | 4.32E-02 |
| Order | Actinomycetales | RSR | Abundance | -3.68E-04 | 1.84E-01 | 6.75 | 13.43 | 1.01E-02 |
| Order | Sphingomonadales | RSR | Abundance | 6.96E-04 | 1.58E-01 | 7.44 | 1.13 | 1.01E-02 |
| Family | Polyangiaceae | RSR | Abundance | 2.19E-02 | 1.37E-01 | 8.01 | 0.08 | 1.81E-02 |
| Family | Sinobacteraceae | RSR | Abundance | -1.48E-03 | 1.67E-01 | 5.8 | 0.05 | 2.96E-02 |
| Family | Sphingomonadaceae | RSR | Abundance | 6.96E-04 | 1.58E-01 | 7.44 | 1.13 | 1.81E-02 |
| Genus | *Minicystis* | RSR | Abundance | -7.91E-04 | -1.32E-02 | 8.74 | 0.00 | 1.58E-02 |

**Table S11.**
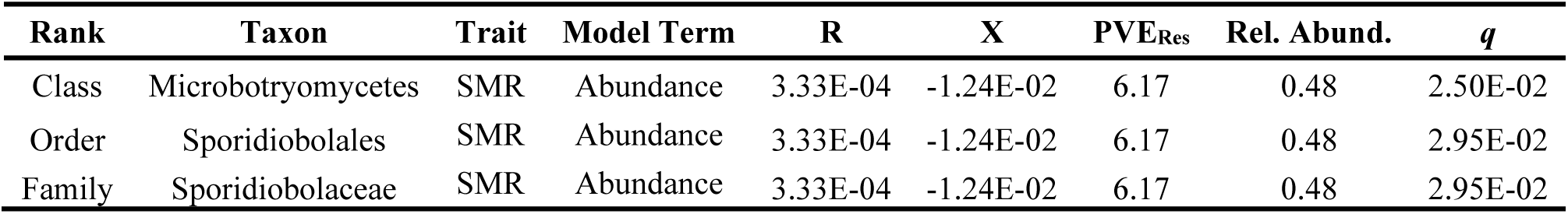
Fungal taxa across taxonomic levels associated with square root transformed shoot mass rate.

| Rank | Taxon | Trait | Model Term | R | X | PVE<sub>Res</sub> | Rel. Abund. | q |
| --- | --- | --- | --- | --- | --- | --- | --- | --- |
| Class | Microbotryomycetes | SMR | Abundance | 3.33E-04 | -1.24E-02 | 6.17 | 0.48 | 2.50E-02 |
| Order | Sporidiobolales | SMR | Abundance | 3.33E-04 | -1.24E-02 | 6.17 | 0.48 | 2.95E-02 |
| Family | Sporidiobolaceae | SMR | Abundance | 3.33E-04 | -1.24E-02 | 6.17 | 0.48 | 2.95E-02 |

